# Mixed-vertebrate pollination traits and pollinators of *Haageocereus acranthus* (Cactaceae) in a lomas desert ecosystem of coastal Peru

**DOI:** 10.64898/2026.01.08.698271

**Authors:** Bernardo Garcia-Simpson, Braulio A. Prado, Juan J. Pellón, Alfonso Valiente-Banuet

## Abstract

Cacti are key components of arid ecosystems and, as predominantly outcrossing plants, rely on animal pollination. Although bee pollination is ancestral, systems supported by birds, moths, bats, and mixed strategies have evolved. Despite the ecological uniqueness and extreme seasonality of the fog-dependent lomas ecosystem of coastal Peru, cacti pollination remains unstudied. This work examines *Haageocereus acranthus*, a common columnar cactus of this ecosystem, in a population where individuals consistently produce either white or pink-red flowers. It was hypothesized that white flowers would be associated with bat pollination and pink-red flowers with hummingbird pollination, reflected in differences in floral morphology, phenology and pollinator visitation.
Year-round monitoring of a population was conducted to characterize flowering phenology. Floral morphology, daily anthesis patterns, and nectar production were quantified and compared between color morphotypes. Floral visitor frequency, behavior and preference were recorded using trail cameras.
Floral phenology, morphology, and nectar characteristics were broadly consistent with vertebrate pollination, showing traits associated primarily with bat pollination but also compatible with hummingbird pollination. These floral traits did not differ between color morphotype. Hummingbirds were the most frequent visitors, followed by bats; yet, neither group showed a preference for a specific flower color.
Findings support a mixed pollination system involving both hummingbirds and bats in an ecosystem where the availability of pollinators could shift over geographic or temporal scales. This mixed strategy could reduce vulnerability to the absence or decline of specific pollinator groups, ensuring consistent reproductive success.

**Key message:** In a lomas desert ecosystem of coastal Peru, the columnar cactus *Haageocereus acranthus* shows traits and interactions consistent with a mixed-vertebrate pollination system (hummingbirds and bats), with no detectable differences between flower color morphotypes in either traits or interactions.

## Introduction

Pollination has played a central role in angiosperm interactions and diversification (Vamosi & Vamosi 2010; Ballesteros-Mejia *et al*. 2016), reflecting the evolutionary innovations of flowers in response to different ecological contexts (Armbruster 2017; Opedal 2019). Species sharing pollinators have usually been under similar selective pressures, converging on suites of floral traits known as pollination syndromes (Faegri & van der Pijl 1979; Fenster *et al*. 2004; Dellinger 2020). These have long been considered a principle governing plant-pollinator interactions and used to infer functional pollinator groups from flower traits (Proctor *et al*. 1996; Fenster *et al*. 2004). However, increasing evidence supports that these patterns are more complex than the traditional syndromes hypothesis suggests (Ollerton *et al*. 2009; Rosas-Guerrero *et al*. 2014; Dellinger 2020), since pollinator assemblages can be highly variable across a species range (Thompson 2002), and interactions often show geographical or temporal changes (Olesen & Jordano 2002; Burkle & Alarcon 2011). Moreover, intraspecific variation in floral traits may cause deviations from classic syndromes, as different variants within a species can rely on distinct pollinator groups (e.g., Schlumpberger *et al*. 2009; Cardona *et al*. 2020; Wenzell *et al*. 2025). This can result in local adaptation, divergence in pollination strategies, and potentially lead to speciation (Kay & Sargent 2009).

Cacti are distributed across the Americas and constitute crucial components of arid and semi-arid ecosystems (Fleming *et al*. 2001; Fleming & Valiente-Banuet 2002). They strongly depend on animals for pollination (Subfamily Cactoideae; Pimienta-Barrios & del Castillo 2002; Mandujano *et al*. 2010), with outcrossing favored by high inbreeding depression or self-incompatibility in many species; though the breeding system remains uncharacterized for the vast majority of the family. Floral adaptations are mostly consistent with classic pollination syndromes (Grant & Grant 1979; Rowley 1980; Schlumpberger 2011). While bee pollination is ancestral in cacti, specialized systems for bird, moth, and bat pollination have evolved (Schlumpberger 2011; Lendel 2013; Gorostiague *et al*. 2023). Also, evidence supports the existence of mixed pollination strategies in which a species relies on more than one pollinator guild; such as bat-hummingbird in Sahley (1996) and Bustamante *et al*. (2010); and hawkmoth-bee in Walter (2010).

Peru is a cacti species-rich country with ∼250 spp. distributed mainly in coastal and Andean ecosystems (Ulloa *et al*. 2004; Arakaki *et al*. 2006). However, although Peruvian cacti have received attention at taxonomic, distributional and genetic levels (e.g., Ostolaza 1996, 2014; Arakaki *et al*. 2006, 2007, 2021; Calderón *et al*. 2007), scarce effort has been made to understand their ecological interactions (e.g., Sahley 1996; Novoa *et al*. 2005, 2022; Ceroni *et al*., 2007). This gap is more relevant in the context of the Peruvian coastal desert: a hyper arid yet humid environment shaped by the interaction of the Andes, the Humboldt Current and the South Pacific High anticyclone (Reynel *et al*. 2013). Within this landscape, winter fogs interact with coastal mountains to generate the lomas: highly seasonal patches of vegetation dominated by annual plants and a subset of perennial species, restricted almost entirely to Peruvian territory (Rundel *et al*. 1991; Moat *et al*. 2021). Here, cacti are among the few perennials of a strongly seasonal ecosystem, and their flowering and fruiting potentially constitute pulse of resources during periods when most vegetation is absent (e.g., Nobel & Loik 1999; Fleming & Valiente-Banuet 2002).

*Haageocereus acranthus* (Vaupel) Backeb. (Tribe Trichocereeae) is a common columnar cactus in the river valleys and lomas of the central coast of Peru. Its flowers are relatively large, robust, funnel-to tubular- shaped, single-opening, nocturnal, white and nectar-abundant; traits putatively associated with bat or sphingid moth pollination, based on classic syndromes (Faegri & van der Pijl 1979; Calderón *et al*. 2007). However, marked intraspecific variation has been documented in flower size, shape, pigmentation (white to pink-red), and extended anthesis times spanning crepuscular and matutinal periods; suggesting the involvement of additional pollinator groups such as hummingbirds. Visitation by *Rodophis vesper* (Oasis hummingbird) and *Platalina genovensium* (Peruvian long-tongued bat) have been reported (Grillo & Arana 2016; Maguiña & Amanzo 2016), and empirical evidence supports a mixed bat-hummingbird system in morphologically similar, and closely related, *Weberbauerocereus weberbaueri* on the southern Peruvian coast (Sahley 1996).

No detailed pollination study has been conducted for *H. acranthus*. This represents a significant gap, not only for the species itself, but for understanding pollination dynamics of cacti in the unique ecological context of the Peruvian coastal desert. Lomas are among the most singular vegetation formations, yet cactus pollination within them remains entirely undocumented. Addressing this constitutes a first step toward assessing the functional role of cacti as resource providers in the lomas, and toward placing Peruvian species within the broader picture of pollination strategies described for Cactaceae.

This study examined the floral morphology, phenology, and floral visitor assemblage of *H. acranthus* in a lomas ecosystem of the Peruvian central desert coast, in a population exhibiting two distinct flower color variants: white and pink-red, each expressed by different individuals. Our goal was to determine the putative pollination syndrome of this species, based on floral traits, and evaluate whether it is supported by empirical evidence of floral visitation and visitor behavior. We hypothesized that *H. acranthus* would display traits consistent with vertebrate pollination, and that bats and hummingbirds would be the most frequent visitors, exhibiting behavior compatible with an effective pollen transfer (i.e., contacting reproductive structures). Based on classic pollination syndromes, we further predicted that the two color morphotypes would differ in their association with specific guilds: white flowers would be more closely linked to bats, while pink-red flowers would show a tendency toward hummingbirds. We anticipated that these flower types would occupy distinct regions of floral morphological space and exhibit different phenological patterns, with pink-red flowers exhibiting a broader anthesis and nectar secretion period extending into diurnal hours, and consequently attracting hummingbirds at higher relative frequencies than white flowers.

## Materials and Methods

### Study area and species

This study was conducted in Cardal (Pachacamac, Lima, Peru) (12°11’S, 76°50’W; 215 m.a.s.l.), a lomas cactus locality previously documented by Ostolaza (1996) and Calderón *et al*. (2007) (Fig. 1A). The lomas are seasonal ecosystems driven by winter coastal fogs, restricted mainly to Peru and Chile, and characterized by a wet season (May–Oct; 13–16°C, 95–100% relative humidity) that promotes the growth of stational herbs, and a dry season (Nov–Apr; 20–25°C, 80–90% relative humidity) during which most herbaceous vegetation dies back, leaving only perennial xerophytes (Dillon et al. 2011). Fieldwork was conducted from January 2022 to March 2023.

**Figure 1.**
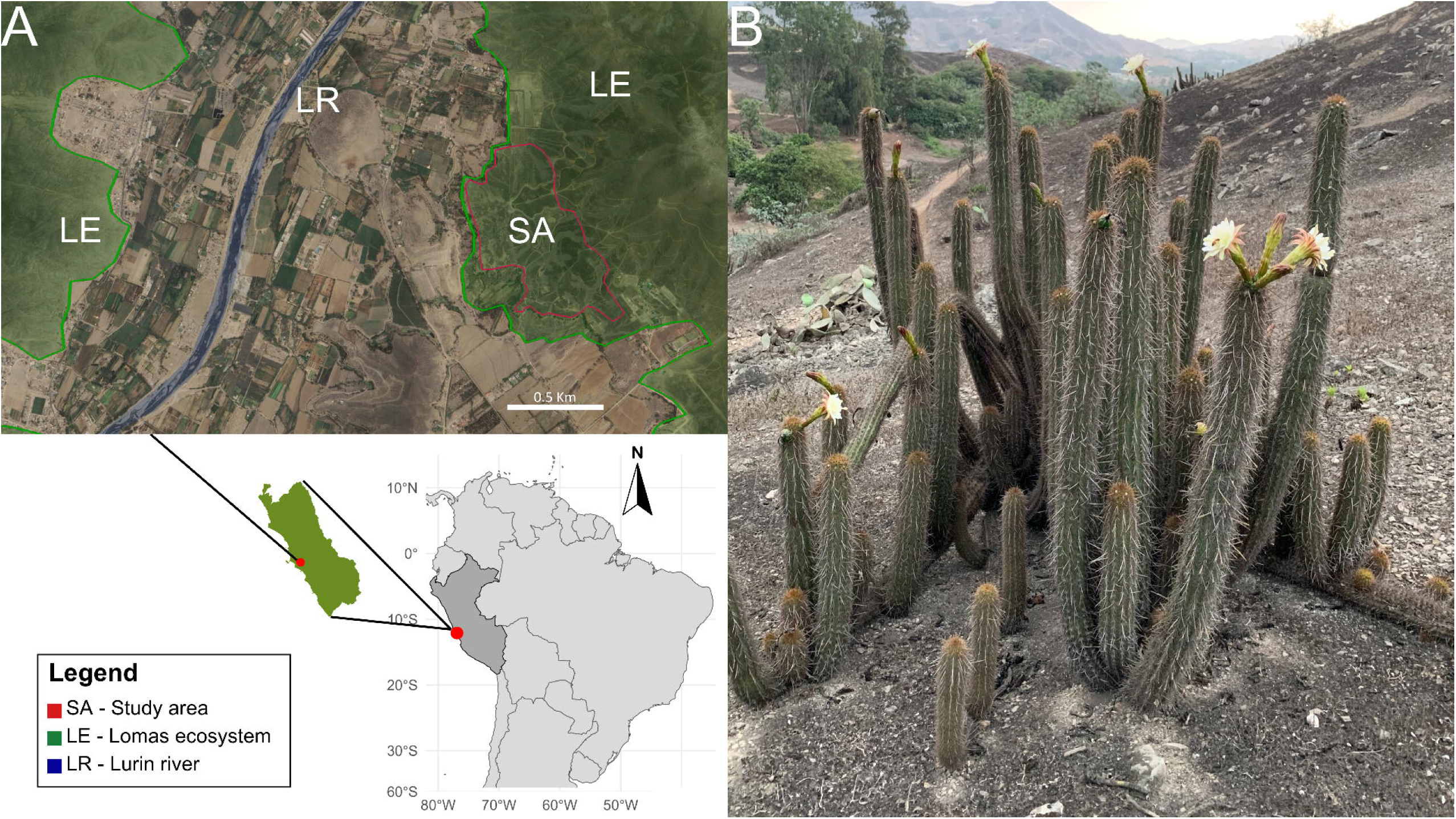
**(A)** Geographic location and composition of the study area in Peru, showing the study area (SA) within the lomas ecosystem (LE), near the Lurin River (LR), and surrounded by agricultural and semi-urban land. **(B)** Individual of *H. acranthus* in the study area.

*H. acranthus* is a columnar cactus up to 1.5 m tall, with multiple vertical stems branching at the base and flowers often growing from areoles at the branch apex (Calderón *et al*. 2007) (Fig. 1B). Aside from observations of flowering from November to January, no detailed phenological information is available (Calderón *et al*. 2007; Maguiña & Amanzo 2016). Two flower types are present in this area: common white flowers of *H. acranthus* subsp. *acranthus* and pink-red flowers of *H. acranthus* var. *olowinskianus* f. *rubriflorior* (Ostolaza 1996), currently a synonym of *H. acranthus* subsp. *acranthus* (Calderón *et al*. 2007; POWO 2024). Flower types are hereafter referred to as white and pink-red floral morphs. Although *H. acranthus* dominates, two other cacti co-occur at much lower densities: *H. pseudomelanostele* (Werderm. & Backeb.) Backeb., which is similar in morphology and phenology to *H. acranthus*; and *Loxanthocereus acanthurus* (Vaupel) Backeb., which exhibits hummingbird pollination traits and flowers at a different time of the year (B. Garcia-Simpson, pers. obs.).

### Annual phenology

We monthly monitored 30 tagged plants throughout 2022, recording the number of floral buds, open flowers and fruits (firm and green for unripe, red and for ripe) per individual. Due to field limitations, monitoring was conducted exclusively on individuals with white flowers. However, observations from recurrent prior and subsequent visits to the study area (years 2021 to 2025; B. Garcia-Simpson, pers. obs.) suggest that the annual phenological pattern of the white morphotype documented here is representative of the broader population.

### Floral traits – morphology

We collected 30 fully open flowers (15 per floral morph), each from a different individual (30 individuals in total). Longitudinal cut sections were photographed in the field with an iPhone XR camera (12MP, f/1.8 aperture, 26mm focal length) (Apple, California, USA) against a black background with a reference scale. Total length (TL), perianth width (PW), tube length (TuL), tube width (TuW), stigma exsertion (StiE), stamen exsertion (StaE), ovary length (OL), ovary width (OW), nectar chamber length (NL), and nectar chamber width (NW) (Fig. 2A–D; terminology modified from Nassar *et al*. 1997) were measured from photos using ImageJ v.1.54f (Schneider *et al*. 2012).

**Figure 2.**
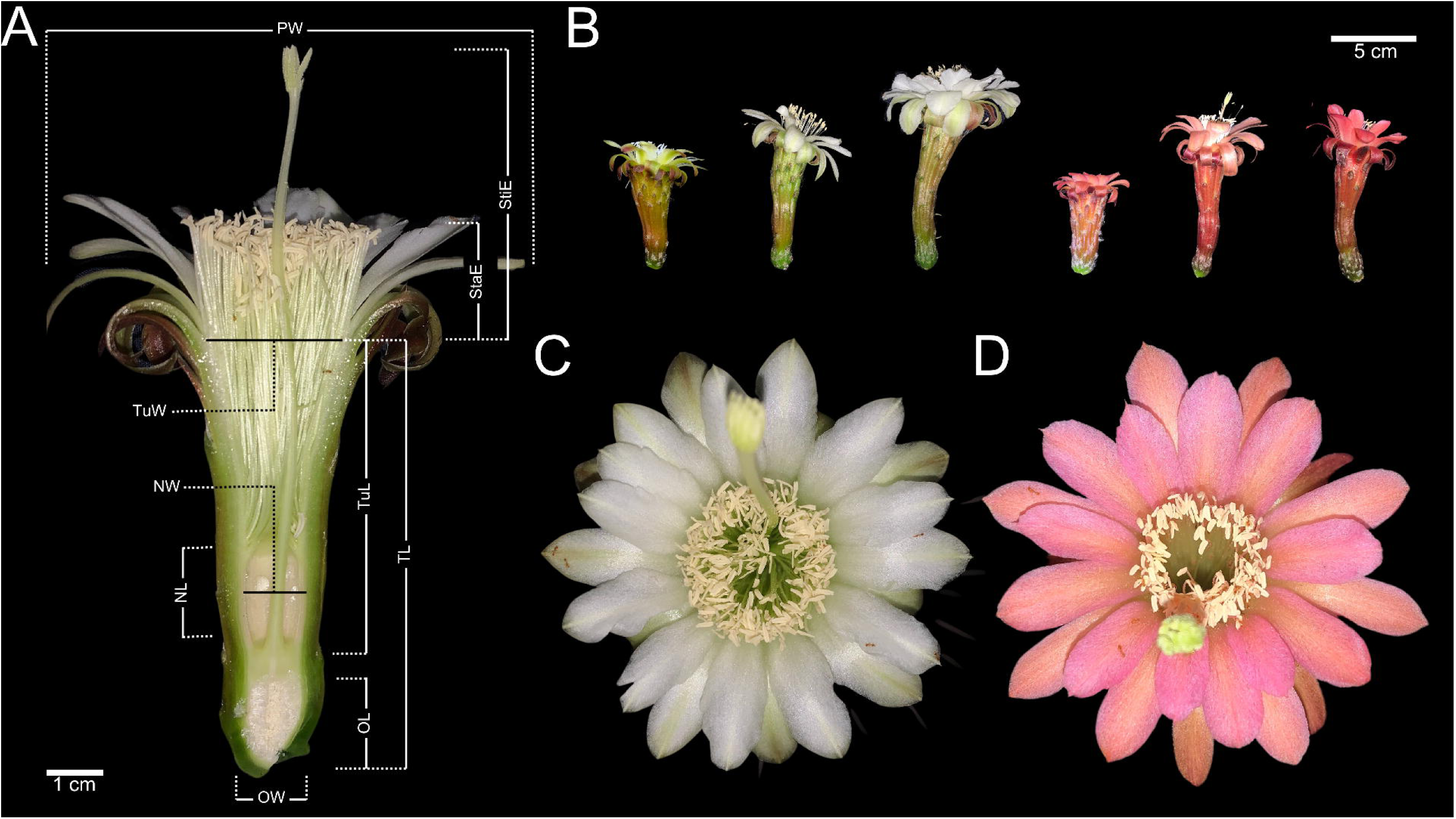
(A) Longitudinal cut of *H. acranthus* flowers indicating measured morphometric variables. (B) Size and shape variation in flowers of both morphotypes. Frontal view showing perianth color and reproductive structures of (C) white and (D) pink-red flower morphotypes.

### Floral traits **–** daily phenology

To characterize flower aperture, we measured the perianth width of 23 flowers (11 white, 12 pink-red) from different individuals (23 individuals in total) at eight stages of anthesis (15:00, 17:00, 19:00, 23:00, 03:00, 05:00, 07:00, 09:00) using a 0.05 mm resolution mechanical caliper (Uyustools, Hangzhou, China). Nectar volume, sugar concentration, and energy content were assessed at five anthesis stages (15:00, 19:00, 23:00, 03:00, 06:00) using bagged flowers from different individuals. Nectar volume was measured in 34 flowers (18 white, 16 pink-red), and sugar content was evaluated from a subset of 17 flowers (10 white, 7 pink-red). Nectar volume was quantified, following Ibarra-Cerdeña *et al*. (2005), by extracting all possible nectar using a capillary tube inserted into the nectar chamber. Total nectar volume was calculated by multiplying the column length by the specified cross-sectional area of the capillary tube. A fraction of the nectar was used to determine sugar concentration with a handheld refractometer (BRIX30) with automatic temperature compensation; readings were expressed as sucrose percentage following Dafni (1992) and energy supply was estimated using the following formula:

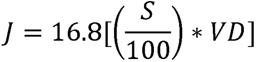

where J is energy in joules, S is the sugar percentage, V is nectar volume in microliters, and D is the density of sucrose at the observed concentration (Dafni 1992; Ibarra-Cerdeña *et al*. 2005).

### Frequency and behavior of floral visitors

Flowers were monitored using two Bushnell 24MP Core Low Glow Trail Cameras (Bushnell Corporation, Kansas, USA) placed 1–2.5 meters to record interactions with vertebrate and large invertebrate visitors, during day and night. A total of 22 (11 white, 11 pink-red) flowers from different individuals were observed over 11 non-consecutive days during the flowering season (Sep 2022–Mar 2023), totaling 378 hours (∼17 hours per flower). Monitoring started in the early afternoon and continued until mid-morning the following day. Following Ibarra-Cerdeña *et al*. (2005), visitor species and number of legitimate visits (i.e., involving contact with reproductive structures) were recorded through complete visual review of all photos and videos. Additional visitors were documented through direct observation.

### Preliminary pollinator exclusion experiments

To evaluate how the exclusion of specific pollinator groups influenced fruit initiation, we implemented five treatments: (i) control, with flowers left fully exposed; (ii) bat exclusion, covering flowers at night with a cylindrical metallic mesh; (iii) hummingbird exclusion, applying the same mesh only during the daytime; (iv) nocturnal exclusion, fully bagging flowers at night to prevent all nocturnal visitors; and (v) diurnal exclusion, fully bagging flowers during the daytime to block diurnal visitors. We monitored each flower over the following days to record fruit initiation. Due to fieldwork constraints, sample sizes were limited to 12 flowers per morph in the control treatment and six flowers per morph in each exclusion treatment. Results are presented descriptively as preliminary evidence, without formal statistical analyses.

### Data analyses

All statistical analyses were conducted in R v.4.4.1 (R Core Team 2024) using RStudio (Posit Team 2024). For all analyses, statistical significance was assessed at α = 0.05.

To evaluate the effect of flower morph (white vs. pink-red) on morphometric variables, we used a MANOVA (e.g., Pérez-Barrales *et al*. 2007). Variables were standardized (mean = 0, standard deviation = 1) for comparability. To avoid multicollinearity, we examined correlations among morphometric variables: total length, tube length, and nectar chamber length were highly correlated (Pearson’s r > 0.8), therefore, we excluded tube length and nectar chamber length from the test. The analysis was performed with the *manova* function.

To examine the effects of time after midday (hours) and flower morphotype (white vs pink-red) on flower aperture (measured as perianth width), nectar volume, nectar sugar concentration, and nectar energetic content, we fitted Generalized Linear Mixed-Effects Models (GLMMs). In the absence of specific hypotheses regarding differences among morphotypes (e.g., response magnitude or peak times), we opted for an exploratory model selection approach by generating a set of biologically plausible candidate models and selecting the best fit.

Model selection was conducted separately for each response variable: candidate models were built using the *glmmTMB* package (Brooks *et al*. 2017), with Flower ID included as a random effect to account for individual variability. Time after midday (T) and flower morphotype (M) were included as fixed effects, testing first-, second-, and third-order orthogonal polynomial terms for time to capture potential curvilinear relationships. Fixed effects combination in candidate models included: T, T + M, T * M, T^2^, T^2^ + M, T^2^ * M, T^3^, T^3^ + M, T^3^ * M. A model including a polynomial term of degree *n*, it inherently incorporated all lower-order terms (e.g., a T³ model also included T and T²). ‘+’ indicates additive effects, while ‘*’ denotes additive and interaction effects. Flower aperture and sugar content were modeled using a Gaussian distribution, while nectar volume and energetic content were modeled with a Tweedie distribution, chosen for its suitability in handling continuous data with a high proportion of zeros. Model diagnostics; presence of over/under-dispersion, outliers, and zero inflation, were assessed using the *DHARMa* package (Hartig 2022). Model selection indices were computed using *compare_performance* from the *performance* package (Lüdecke *et al*. 2021) and *model.sel* from the *MuMIn* package (Barton 2024). Final selection was based on the Akaike Information Criterion corrected for small sample sizes (AICc), with models differing by ≥ 2 AICc units considered significantly better fits (Burnham & Anderson 2002). In addition, to assess the stability of model estimates, we conducted a cluster bootstrap procedure (Field & Welsh 2007), resampling individual flowers with replacement, retaining all associated repeated measurements, and refitting the selected model to each resampled dataset (1000 iterations). The resulting distribution of estimates across iterations was compared against the original model estimates and 95% bootstrap confidence intervals.

To visualize the temporal activity patterns of main floral visitors, we used a kernel density estimation (KDE) with the *density* function (bandwidth = 1). Densities were scaled by the total number of visits to make activity patterns visually comparable across visitor groups with differing number of visits.

To assess if the number of visits varied among floral visitors and flower types, we built Generalized Linear Mixed-effects Models (GLMMs) with a Poisson distribution (log-link), as visit counts consist of non-negative integers. Flower ID was considered as a random effect, while visitor type (bat, hummingbird or sphingid) and flower morphotype were considered as fixed effects. We evaluated three models including: visitor type (V), visitor type and flower morphotype (V + M), and visitor type, flower morphotype, and their interaction (V * M). To account for differences in evaluation time among individual flowers, all models included the log-scaled evaluation time of each flower as an offset variable. Model fitting, diagnostics, and selection were performed as previously described. Based on the best-supported model, post-hoc pairwise comparisons of visitation frequencies among floral visitors were conducted using the *emmeans* package (Lenth 2018). Pairwise differences were evaluated with the *pairs* function, applying a Tukey adjustment to account for multiple comparisons.

## Results

### Annual phenology

*Haageocereus acranthus* from Cardal displayed a single flowering peak during the year. Bud and flower production spanned from January to April. From May to September, plants remained mostly vegetative. Bud production resumed in October, peaking in November, resulting in a high number of open flowers in December. Fruit production followed this pattern, with a slight delay due to the sequential nature of these reproductive stages. Overall, reproduction occurred during the dry season (Nov–Apr), while the vegetative phase in the humid season (May–Oct) (Fig. 3A–B; Table S1).

**Figure 3.**
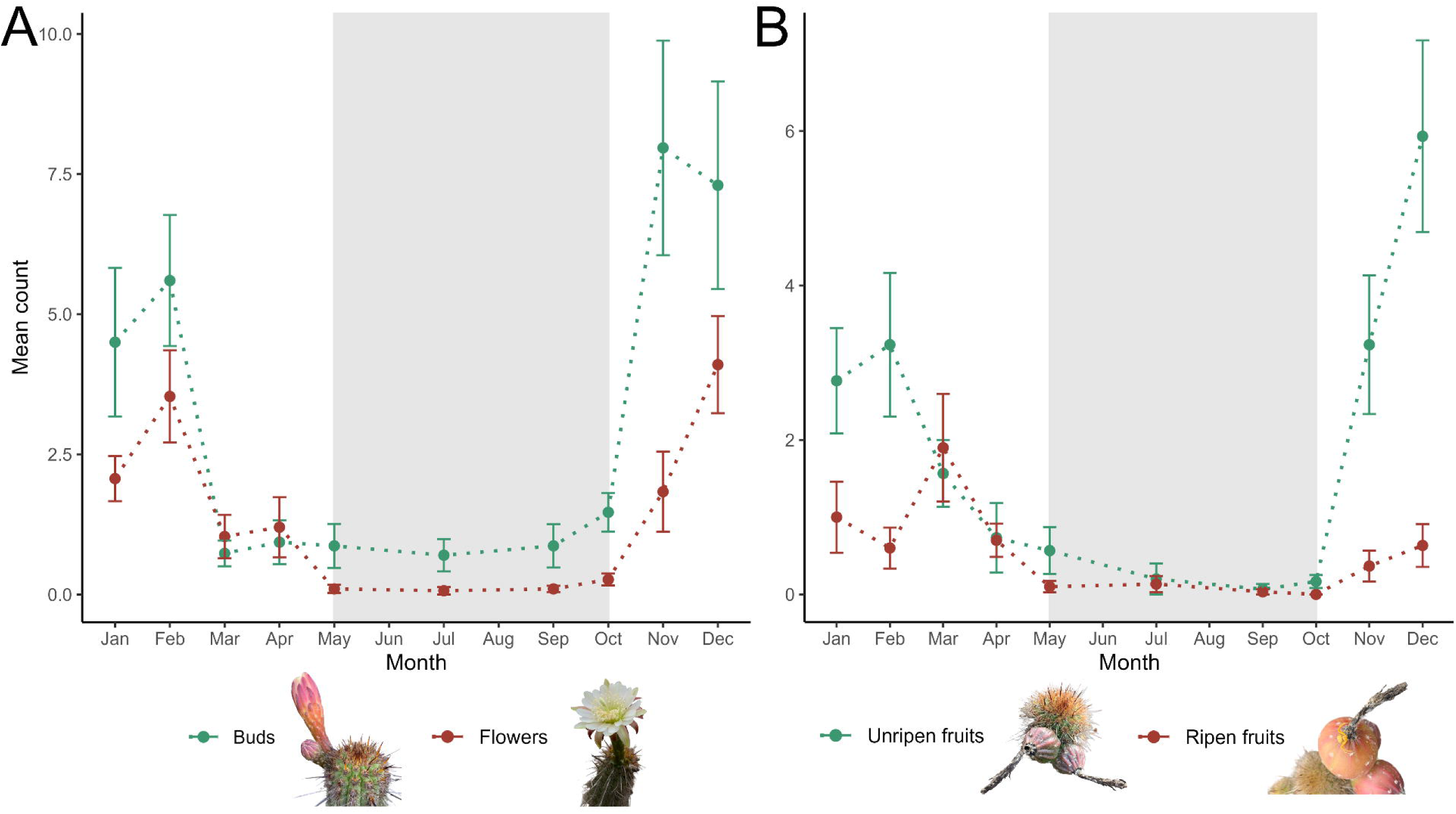
Monthly mean counts of (A) floral buds and flowers, and (B) unripe and ripe fruits observed throughout 2022. Vertical bars indicate the standard error of the mean. Shaded gray areas correspond to the humid Lomas season.

### Floral traits – morphology

*Haageocereus acranthus* flowers were funnelform to tubular in shape and radially symmetric, but occasionally exhibited a slight degree of zygomorphy (Fig. 2B). The flower tube exhibited a constriction above the nectar chamber, which could limit access to nectar. Perianth color remained consistent within individuals, with each plant producing exclusively white or pink-red flowers (Fig. 2C–D). Although some intra-morph color variation was observed (white flowers ranged from whitish-green to pure white; pink-red flowers from carmine-pink to reddish), these were clearly distinct with no intermediate colors. Overall morphometry of the two floral morphs did not differ (Figure S1, Table S2), as supported by the MANOVA results (Pillai’s Trace = 0.221, *F* (8, 23) = 0.813, *P* = 0.598).

### Floral traits – daily phenology

Unless otherwise indicated, model estimates are reported with 95% confidence intervals (mean ± 95% CI).

#### Flower aperture

The top-ranked model included time up to the cubic term (T, T², T³) and its respective interactions with morphotype (AICc = 467.29, weight = 0.560). The second-best model additionally included the main effect of morphotype (ΔAICc = 2.31, weight = 0.176) (Table S3). Diagnostics were overall acceptable, and bootstrapped coefficient estimates indicated stable and reliable fixed effects for the selected model (Fig. S2–S3). The linear effect of time was weak, whereas the quadratic component had a strong effect, reflecting the clear parabolic pattern in flower aperture over time, consistent with the expected non-linear dynamics of anthesis. The cubic term introduced a slight degree of asymmetry into this pattern, whereby the decline in aperture following the maximum was more gradual and prolonged than the initial opening phase. Both floral morphs reached maximum aperture around 23:00 (white: 6.51 ± 0.3 cm, pink-red: 5.9 ± 0.3 cm). An interaction with morphotype was also detected, with the white morph initiating aperture slightly earlier (white: 14:34, pink-red: 12:81) and reaching a marginally higher maximum aperture than the pink-red morph (Fig. 4A, Table 1A). However, the weak effect size associated morphotype interactions suggest that, while statistically detectable, their biological relevance may be limited. Styles and stigmas remained turgid throughout anthesis.

**Figure 4.**
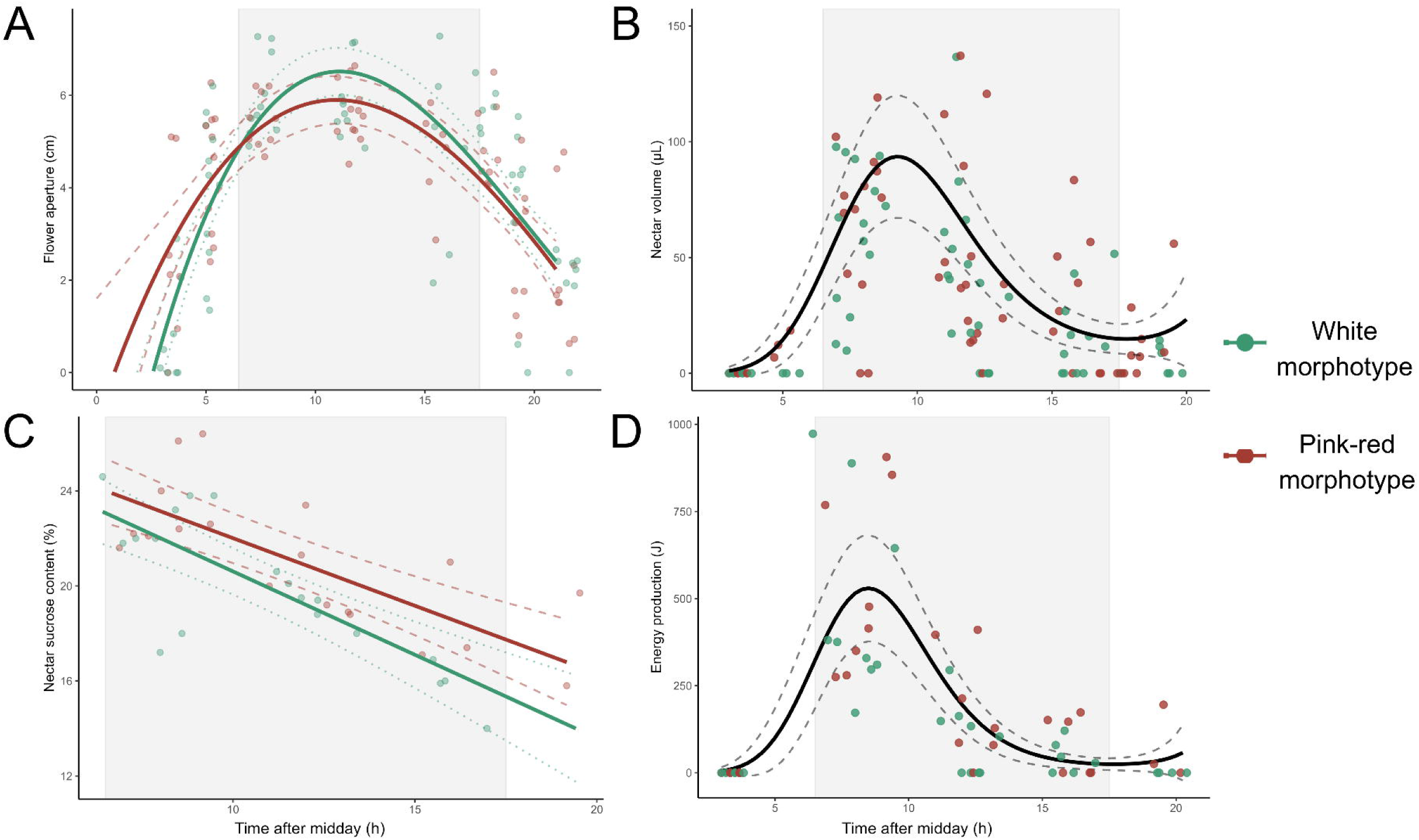
Temporal variation in (A) flower aperture, (B) nectar volume, (C) nectar sucrose content, and (D) nectar energy production as a function of time after midday. Solid lines represent model predictions, and dotted and dashed lines indicate 95% confidence intervals. In panels A and C the model includes the effect of morphotype, providing separate estimates and confidence intervals for each morph. Shaded gray areas correspond to the nocturnal period.

**Table 1.** Summary of the best-fitting models for each response variable: (A) flower aperture, (B) nectar volume, (C) nectar sugar concentration, (D) nectar energy content, and (E) visitation frequency. Fixed effects are expressed as estimates with their standard errors (SE), Z-values (*Z*), and p-values (*P*). Estimates are relative to white flowers for morphotype and to sphingid moths (*Manduca* sp.) for floral visitors. ‘Morph’ and ‘T’ represent the effect of morphotype and time after midday, respectively; ‘:’ denotes interaction among terms.

| Response variable | Fixed effect | Estimate | SE | Z | P |
| --- | --- | --- | --- | --- | --- |
| **(A) Flower aperture** | Intercept | 4.11 | 0.17 | 24.88 | <0.001 |
|  | T | -0.97 | 1.59 | -0.61 | 0.540 |
|  | T<sup>2</sup> | -18.83 | 1.58 | -11.93 | <0.001 |
|  | T<sup>3</sup> | 4.34 | 1.54 | 2.83 | <0.005 |
|  | T : Morph red-pink | -4.32 | 2.26 | -1.91 | 0.056 |
|  | T<sup>2</sup> : Morph red-pink | 4.89 | 2.24 | 2.18 | 0.029 |
|  | T<sup>3</sup> : Morph red-pink | -2.61 | 2.25 | -1.16 | 0.247 |
| **(B) Nectar volume** | Intercept | 3.33 | 0.15 | 22.45 | <0.001 |
|  | T | -0.11 | 2.05 | -0.05 | 0.957 |
|  | T<sup>2</sup> | -9.28 | 1.76 | -5.29 | <0.001 |
|  | T<sup>3</sup> | 8.14 | 1.81 | 4.51 | <0.001 |
| **(C) Nectar sugar content** | Intercept | 19.61 | 0.50 | 38.93 | <0.001 |
|  | T | -15.51 | 2.58 | -6.02 | <0.001 |
|  | Morph red-pink | 1.59 | 0.72 | 2.22 | 0.027 |
|  | T : Morph red-pink | 2.94 | 3.41 | 0.86 | 0.389 |
| **(D) Nectar energy** | Intercept | 4.33 | 0.22 | 19.69 | <0.001 |
|  | T | -1.19 | 2.55 | -0.47 | 0.639 |
|  | T<sup>2</sup> | -7.71 | 2.06 | -3.74 | <0.001 |
|  | T<sup>3</sup> | 8.54 | 1.79 | 4.78 | <0.001 |
| **(E) Visitation frequency** | Intercept | -5.63 | 0.72 | -7.83 | <0.001 |
|  | Visitor *P. genovensium* | 1.20 | 0.47 | 2.59 | 0.01 |
|  | Visitor *R. vesper* | 2.59 | 0.42 | 6.12 | <0.001 |

#### Nectar volume

The best model for nectar volume included time up to the third-order polynomial term but excluded interactions with morphotype and its main effect (AICc = 1025.65, weight = 0.920), Indicating that nectar secretion dynamics were consistent across both floral morphotypes. The interactions and main effect of morphotype were present in models of lower rank (Table S4). Diagnostics were overall acceptable, and bootstrapped coefficient estimates indicated stable and reliable fixed effects for the selected model (Fig. S4-S5). The quadratic effect was significant with a strong effect size, reflecting a clear parabolic pattern in nectar production over time. The additional cubic term introduced asymmetry into this pattern: nectar volume rose at an initially slow but progressively accelerating rate during the late afternoon, reaching a peak of 93.5 ± 13.5 µL at approximately 21:00, after which the decline was considerably slower and more prolonged than the initial rise. This asymmetry resulted in low but detectable quantities of nectar remaining available through the night and into the following morning, before the flower ultimately closed. No significant effect of morphotype was detected, indicating these dynamics were consistent across white and ping-red morphotypes (Fig. 4B, Table 1B).

#### Sugar content

The three best-ranked models showed ΔAICc < 2 among them. Following a parsimony principle, we selected the model including the linear effect of time, the main effect of morphotype and their interaction (AICc = 165.10, weight = 0.152). This model showed equivalent support compared to a more complex one including the second and third-order effect of time (AICc = 163.28, weight = 0.379); a simpler model including only the effect of time showed clear residual structure, thus was not considered (Table S5). Diagnostics were overall acceptable, and bootstrapped coefficient estimates indicated stable and reliable fixed effects for the selected model (Fig. S6-S7). Mean sugar concentration followed a steady decline throughout anthesis from 23.5 ± 0.5% at 18:30 to 15.4 ± 0.8% by 07:40 (Fig. 4C, Table 1C). Although present in the model, the weak effect of the interaction term and main effect of morphotype suggest that the rate of decline was consistent across morphs and that pink-red flowers maintained a slightly higher baseline sugar concentration than white flowers. However, the small effect sizes of these terms suggest that this difference may be of limited biological relevance.

#### Energy content

The best model for nectar energy content included time up to the third-order polynomial term but excluded morphotype effects (AICc = 540.06, weight = 0.738). The second-best model, which added interactions with morphotype, had less support (ΔAICc = 3.05, weight = 0.160). The main effect of morphotype was included in models of lower rank (Table S6). Diagnostics were overall acceptable, and bootstrapped coefficient estimates indicated stable and reliable fixed effects for the selected model (Fig. S8-S9). Energy content closely mirrored nectar volume dynamics, following and asymmetric parabolic pattern driven by strong quadratic and cubic effects. Values rose progressively during the late afternoon, peaking at 529.8 ± 77.8 J around 20:30, after which the decline was considerably slower and more prolonged than the initial rise. The exclusion of morphotype from the best-supported model indicates that these dynamics were consistent across floral morphs (Fig. 4D, Table 1D).

### Frequency and behavior of floral visitors

We recorded 13 floral visitor species (11 invertebrates, 2 vertebrates; Table 2): nine did not contact reproductive structures and are unlikely pollinators, two made occasional contact, and two consistently contacted both anthers and stigmas, qualifying as effective pollinators.Among small visitors, two ant species (Formicidae) frequently visited flowers at night in large numbers (> 20 individuals), harvesting nectar until senescence (Fig. S10A–C). One sap beetle species (*Carpophilus* sp., Nitidulidae) was found inside flowers, likely feeding on pollen or plant tissue, often in older flowers (Fig. S10D). Three spider species (Anyphaenidae, Thomisidae, and Salticidae) used the external zones of floral tubes and perianth as hunting ground (Fig. S10E–F; Fig. S11A–B). Two Diptera species were recorded; one abundant and found inside floral tubes (Phoridae) (Fig. S11C), and one syrphid observed once feeding on pollen. *Apis mellifera* frequently observed collecting pollen but rarely contacting the stigma, except whenin larger groups (> 5 individuals) (Fig. S11D), while thehalictid *Caenohalictus* sp. was also recorded (Fig. S11E–F).

**Table 2.** Summary of registered floral visitors, including visitor group, taxonomic classification, visit type, floral resource utilized, and activity period. ‘Í and ‘Ĺ represent illegitimate and legitimate visits to flowers, respectively, while ‘I/L’ indicates species that showed both visit types across observations.

| Visitor group | Order | Family | Species | Visit type | Floral resource | Activity period |
| --- | --- | --- | --- | --- | --- | --- |
| **Bees** | Hymenoptera | Apidae | *Apis mellifera* | I/L | Pollen | Diurnal |
|  |  | Halictidae | - | I | Pollen | Diurnal |
| **Ants** | Hymenoptera | Formicidae | *Solenopsis* sp. | I | Nectar | Diurnal/Nocturnal |
|  |  | Formicidae | *Linepithema* sp. | I | Nectar | Diurnal/Nocturnal |
| **Flies** | Diptera | Phoridae | - | I | Pollen | Diurnal |
|  |  | Syrphidae | - | I | Pollen | Diurnal |
| **Beetles** | Coleoptera | Nitidulidae | *Carpophilus* sp. | I | Pollen, plant tissue | Diurnal/Nocturnal |
| **Spiders** | Araneae | Salticidae | - | I | Hunting ground | Diurnal/Nocturnal |
|  |  | Anyphaenidae | - | I | Hunting ground | Diurnal/Nocturnal |
|  |  | Thomisidae | - | I | Hunting ground | Diurnal/Nocturnal |
| **Sphingids** | Lepidoptera | Sphingidae | *Manduca* sp. | I/L | Nectar | Nocturnal |
| **Hummingbirds** | Apodiformes | Trochillidae | *Rhodopis vesper* | L | Nectar | Diurnal |
| **Bats** | Chiroptera | Phyllostomidae | *Platalina genovensium* | L | Nectar | Nocturnal |

Among large visitors, the hummingbird *Rhodopis vesper* (Trochilidae) was the most frequent (percentage of total visits: white: 71.4%, pink-red: 81.4%), active in the late afternoon (17:00–19:00) and early morning (05:00) until the end of anthesis (9:00–11:00) (Fig. 5A–B). They hovered or landed on flowers to access nectar, always contacting stamens and stigma. The bat *Platalina genovensium* (Phyllostomidae) was the second most frequent visitor (white: 20.6%, pink-red: 16.3%), active at night (20:00–03:00, peaking at 22:00) (Fig. 5C–D). Unlike *R. vesper*, bats made single contacts to flowers while hovering but never landed on them. Hawkmoth visits (*Manduca* sp., Sphingidae, Lepidoptera) were rare (white: 7.9%, pink-red: 2.3%), recorded at dusk or night, with inconsistent contact with reproductive structures, (Fig. 5E–F, Table S7).

**Figure 5.**
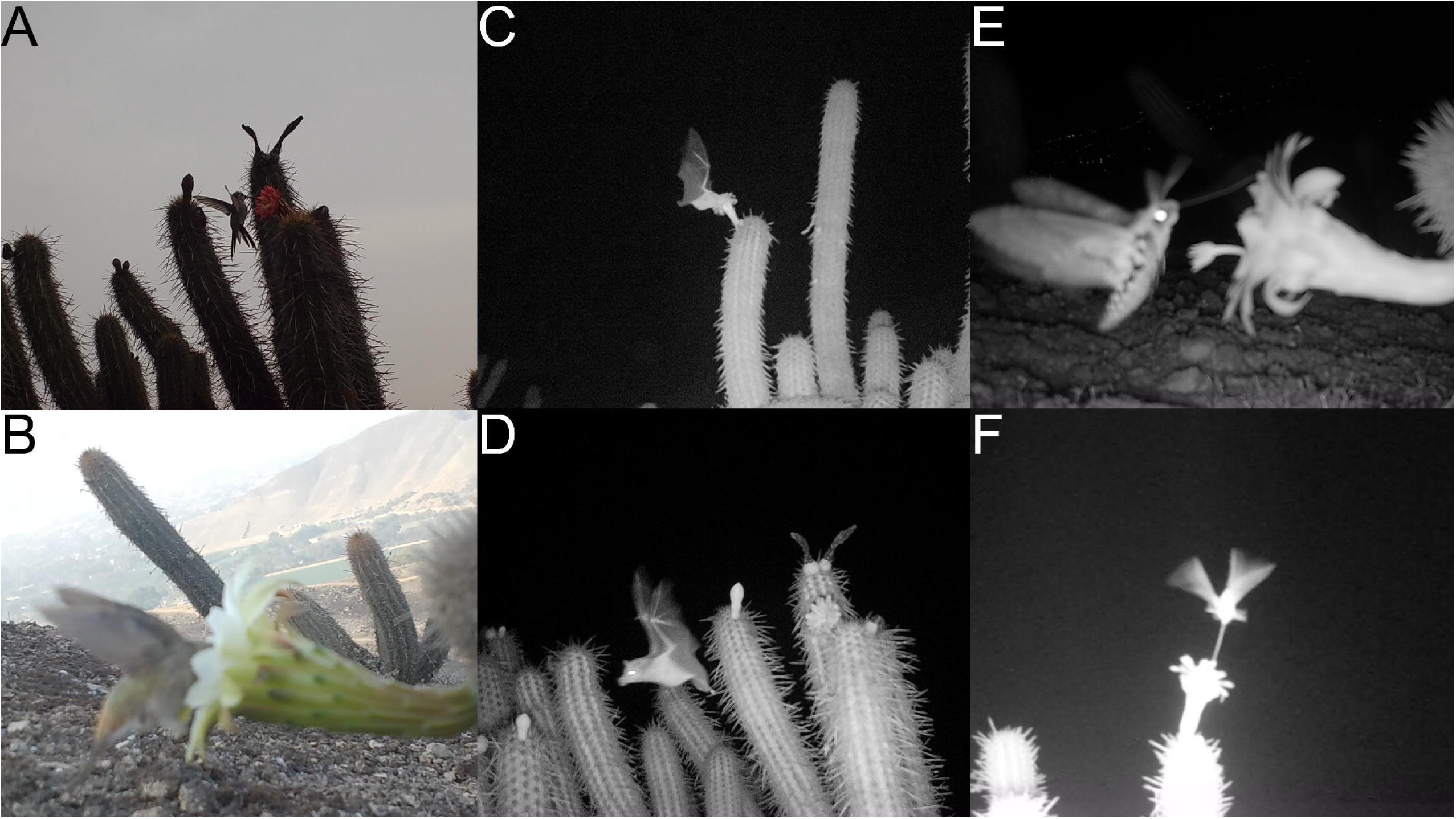
Floral visitors of *H. acranthus*: (A–B) *Rhodopis vesper* (Trochilidae); (C–D) *Platalina genovenisum* (Phyllostomidae); (E–F) *Manduca* sp. (Sphingidae).

**Figure 6.**
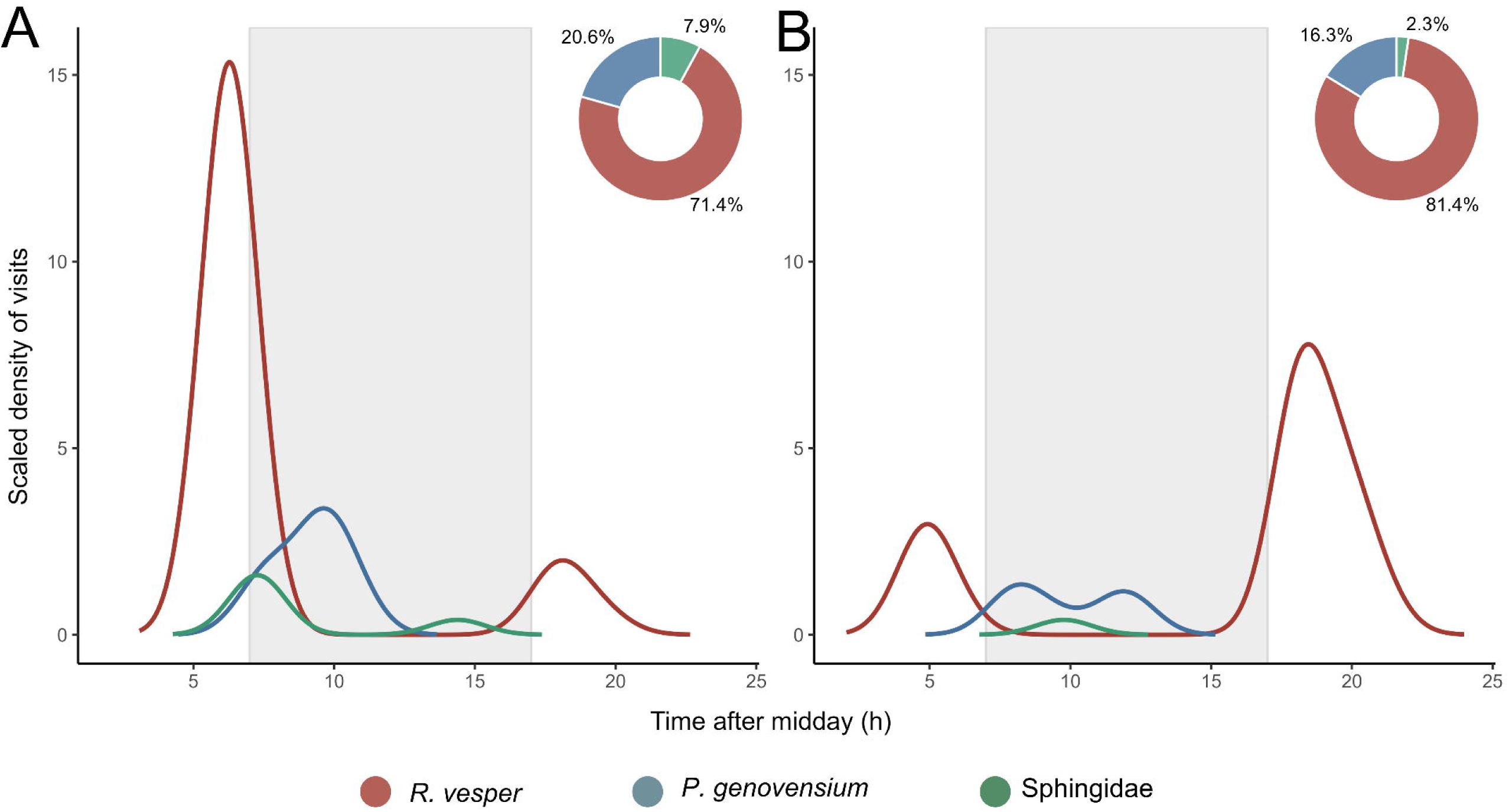
Scaled density of floral visits of *Rhodopis vesper*, *Platalina genovensium*, and sphingid moths over time for (A) white and (B) pink-red flower morphotypes. Ring plots indicate the percentage of total visits contributed by each visitor group for each morphotype. Shaded gray regions indicate the nocturnal period. Shaded gray areas correspond to the nocturnal period.

The best-fitting model included only flower visitor identity (AICc = 209.6, weight = 0.717; Table S8). Diagnostics were overall acceptable, and although bootstrapped coefficient estimates showed wide confidence intervals, mean bootstrap estimates closely matched the reported model coefficients, supporting the reported effects (Fig. S12-S13). Visitation rates differed significantly among visitors: sphingids visited least frequently, significantly less than both *P. genovensium* (estimate = -1.20, SE = 0.465, *P* = 0.0262) and *R. vesper* (estimate = -2.59, SE = 0.423, *P* < 0.001). Additionally, *P. genovensium* exhibited lower visitation rates than *Rhodopis vesper* (estimate = -1.39, SE = 0.250, *P* < 0.001) (Table 1E).

### Preliminary pollinator exclusion experiments

Control flowers showed the highest fruit initiation rates (white: 41.7%; pink-red: 33.3%). Bat exclusion reduced fruit initiation in the white morph but not in the pink-red morph (white: 16.7%; pink-red: 33.3%). Hummingbird exclusion lowered fruit initiation in both morphs (white: 16.7%; pink-red: 16.7%). Nocturnal exclusion produced moderate initiation in the white morph but low initiation in the pink-red morph (white: 33.3%; pink-red: 16.7%), and diurnal exclusion resulted in the same pattern (white: 33.3%; pink-red: 16.7%). In general, fruit initiation was low across all treatments. When morphs were pooled, most treatments showed a fruit initiation rate of 25%, except for the hummingbird-exclusion treatment, which showed 17%. The control flowers exhibited the highest fruit initiation rate at 37.5%.

## Discussion

### Annual phenology

Lomas formations are marked by strong seasonality in floral resources, with most plants reproducing during the humid season (Tovar *et al*., 2018). In contrast, *H. acranthus* flowers during the dry season, a pattern consistent with records from Lachay National Reserve (Maguiña & Amanzo 2016) and other localities (Calderón *et al*. 2007). This recurring pattern across populations of *H. acranthus* suggests that rising temperatures (SENAMHI; Figure S1), and reduced humidity may act as cues for bud development, although this requires further evaluation. Flowering during resource-limited periods positions *H. acranthus* as a potentially key species in this ecosystem, providing critical resources for associated fauna.

### Floral morphology, phenology, and expected pollination syndrome

Morphology of *H. acranthus* reflects a suite of adaptations consistent with other vertebrate-pollinated cacti: its large, sturdy flowers can withstand landings and manipulations by large visitors, while the broad frontal area (∼6.3 cm diameter) likely enhances detection by bats through echolocation, as shown for *Pachycereus* (González-Terrazas *et al*., 2016). Each flower bears 200–400 stamens (Calderón *et al*., 2007), indicating high pollen output and seed set. Comparable bat-pollinated species produce 10□–2×10□ pollen grains per flower (Nassar *et al*., 1997), with *Pilosocereus* reaching ∼500 stamens, ∼2000 ovules, and ∼1100 seeds per fruit (Martins et al., 2020). Together, these traits suggest that *H. acranthus* depends on pollinators with high pollen-carrying capacities within its short floral lifespan (< 24 h).

The floral tube is key for plant-pollinator interactions in cacti (Schlumpberger 2011). In *H. acranthus*, an average tube length of 4.8 cm restricts nectar access to visitors with sufficiently long feeding structures. Some species with shorter structures can still exploit the resource by inserting their head or rostrum into the funnel-shaped tube (e.g., *R. vesper*; Fig. 5B). Tube width influences foraging behavior: wider tubes extend the feeding reach of nectar-feeding bats (Winter & von Helversen 2003; Nicolay & Winter 2006), whereas narrower tubes increase hummingbird handling times (Smith *et al*. 1996). Visitors with feeding structures exceeding ∼6.8 cm (tube length plus stamen exsertion) may obtain nectar without contacting the reproductive organs, thereby acting as nectar robbers (e.g., sphingid moths; Fig. 5F).

Flower phenology confirmed that *H. acranthus* flowers open only once, with peaks in aperture, nectar secretion, and energy availability occurring at night (Fig. 4A–D). Total nectar production, sugar content, and energetic supply were high, comparable to values reported for cacti pollinated by hummingbirds and bats (e.g., Rowley 1980; Grant & Grant 1980; Schlumpberger et al. 2009, Schlumpberger 2011; Albuquerque-Lima *et al*. 2023), suggesting an association with nocturnal vertebrate pollination (Baker 1975; Medel *et al*. 2022 Although pink-red flowers opened slightly earlier than white flowers, nectar characteristics did not differ between morphotypes. Notably, anthesis extended into the late afternoon and morning with small amounts of nectar available, allowing diurnal vertebrates to forage and potentially contribute to pollination (Baker 1961; Miyake & Yahara 1999).

Although pink-red and white pigmentation are traditionally linked to diurnal hummingbird and nocturnal bat/sphingid pollination respectively, our analyses revealed no meaningful differences between white and pink-red floral morphs. Overall, floral traits support primary adaptation to bat pollination, with characteristics also compatible with hummingbird pollination, including diurnal nectar availability and a broad floral tube compatible with hummingbird bill morphology. The relatively short floral tube indicates only limited adaptation to sphingid pollination.

### Empirical evidence of flower visitors and pollinators

Visitors are grouped into three functional categories: (1) ants, sap beetles, flies, and arachnids, which rarely contact reproductive organs and contribute little to no pollination; (2) bees and sphingid moths, which may occasionally transfer pollen but more often act as nectar or pollen thieves; and (3) vertebrates, whose morphology and behavior align most closely with floral traits, making them the expected most effective pollinators.

#### Ants, sap beetles, flies and arachnids

Ants were frequently observed in large groups on flowers but their small size and foraging behavior suggests negligible contributions to pollen transfer (Rico-Gray 1989; Fagua & Ackerman 2011; LeVan *et al*. 2014). *Carpophilus* beetles acted as pollen thieves, feeding on nectar, laying eggs in buds whose larvae consume decaying flowers (Grant & Connell 1979; Miranda-Jácome *et al*. 2021). Dipterans frequently visited but rarely contacted reproductive organs due to their small size, making them ineffective pollinators (Rowley 1980; McIntosh 2005; Schlumpberger *et al*., 2009). Spiders were exclusively predatory visitors, using flowers as hunting grounds (Su *et al*. 2020), though occasional resource use has been reported (e.g., Nelson 2023).

#### Bees and sphingid moths

Bees are frequent visitors due to their pollen-collecting behavior (Westerkamp 1996; Thorp 2000) and can be effective cross-pollinators when morphologically compatible (Armbruster *et al*. 1989; Solís□Montero & Vallejo□Marín 2017). In *H. acranthus*, however, the large flowers and exserted stigmas limit bee-stigma contact, likely reducing their role as efficient pollen vectors. Studies on tropical cacti with similar floral traits and growth habit suggest that the contribution of bees and other small insects to pollination is low or null (e.g., bees, flies, and butterflies on *Weberbauerocereus* in Sahley 1996; *A. mellifera* on *Pilosocereus* in Rivera-Marchand & Ackerman 2006; *Xylocopa grisescens* on *Pilosocereus* in Rocha *et al*. 2019; *A. mellifera* on *Cipocereus* in Martins *et al*. 2020). though, in temperate zones bees often act as effective pollinators, even in bats or moth syndrome species, reflecting more generalized pollination strategies (Valiente-Banuet et al. 1996; Fleming *et al*. 2001).

The traits also suggest sphingid moth pollination (e.g., white pigmentation, nocturnal anthesis, nectar rewards). But sphingids accounted for less than 10% of visits and were observed both landing on flowers or accessing nectar without stamens or stigma contact. This has been previously documented, as it allows feeding from relatively short-tubed flowers of while avoiding approaches that increase predation risk (Wasserthal 2001). By contrast, species adapted to sphingid pollination usually possess much longer floral tubes (e.g., > 10 cm in Grant & Grant 1979; > 15 cm in Silva & Sazima 1995; > 15 cm in Schlumpberger *et al*. 2009; ∼24 cm in Albuquerque-Lima *et al*. 2023). Thus, the role of hawkmoths as pollinators of *H. acranthus* results unlikely.

#### Bats and hummingbirds

Hummingbird foraging overlapped with anthesis only during crepuscular and early morning hours, yet their visits accounted for a major proportion of total observations. The high frequency of *R. vesper* visits highlights its potential role as a pollinator, even though nectar availability at these times was often low or undetectable. This likely reflects repeated nectar extraction throughout anthesis, meaning morning values cannot rule out persistence in flowers unvisited during the night.

Hummingbird-pollinated cacti typically display red to orange, less robust, zygomorphic, and tubular flowers (e.g., *Loxanthocereus*, *Borzicactus*, *Matucana*) (Grant & Grant 1979; Rowley 1980; Schlumpberger 2011). Although *H. acranthus* does not fully match this bauplan, *R. vesper* shows secondary compatibility to access nectar while contacting reproductive structures, with visitation rates not differing between morphotypes. The frequent and effective visits of *R. vesper* support its role as a compatible pollinator, consistent with reports of this species visiting Peruvian cacti lacking typical ornithophilous traits (Sahley 1996; Novoa *et al*. 2022).

The synchronization of *H. acranthus* phenology with the nocturnal activity of *P. genovensium*, coinciding with peak flower aperture and nectar availability is consistent with bat pollination (e.g., Valiente-Banuet *et al*. 1996; Nassar *et al*. 1997; Ibarra-Cerdeña *et al*. 2005; Rocha *et al*. 2019; Martins *et al*. 2020; Albuquerque-Lima *et al*. 2023). Morphologically, *P. genovensium* is well-suited to exploit these flowers, as its long feeding structures match the floral tube. Observations of individuals carrying *Haageocereus* pollen (Maguiña & Amanzo 2016), along with its status as a cactus specialist (Sahley & Baraybar 1996), further support its role as a pollinator. Although bats were not the most frequent visitors, they exhibited the greatest morphological fit to the flowers, suggesting higher pollen transfer efficiency per visit compared to hummingbirds (Muchhala & Thomson 2010). Thus, *P. genovensium* may represent the most effective pollinator of *H. acranthus*, as *Leptonycteris yerbabuenae* does for several columnar cacti in North America below 22° latitude (e.g., Tremlett *et al*. 2020).

Preliminary pollination exclusion experiments in other lomas (Lachay, northern Lima) suggest that *H. acranthus* is primarily pollinated at night (Grillo & Arana 2016). In Lachay, hummingbirds visited flowers frequently, but bat activity was slightly higher. Combined with our observations, this evidence indicates that *H. acranthus* is mainly adapted to bat pollination, though the predominance of chiropterophily may depend on the presence of *P. genovensium*. This bat species is highly dependent on cactus flowers and intact arid habitats, and its distribution is restricted to well-preserved areas that are rapidly declining in Lima (Ossa et al. 2020; Lambert 2021). Accordingly, *P. genovensium* has not been reported from urban environments, where only generalist bats persist by feeding on exotic plants (Pellón *et al*. 2021). The peri-urban setting of our study site, surrounded by towns and agriculture, contrasts with Lachay, a conserved National Protected Area (Dourojeanni 2018), likely explaining the greater prevalence of hummingbird visitation at Cardal. In addition, our preliminary exclusion experiments showed that, although fruit production been generally low, all exclusion treatments tended to produce equal or lower fruit initiation than open flowers, suggesting that both diurnal and nocturnal pollinators contribute to fruit set. Together, these patterns support that the mixed-vertebrate pollination system of *H. acranthus* may facilitate its persistence even where its ideal bat pollinators decline or are absent.

Our findings partially align with those of Sahley (1996) for *Weberbauerocereus weberbaueri* in Arequipa (southern Peru). This closely related species (Tribe Trichocereeae) shares similar morphological and phenological floral traits and is also primarily visited by *P. genovensium* and *R. vesper* (although in *W. weberbaueri* the wine-colored flowers are strongly zygomorphic, unlike in *H. acranthus)*. Sahley (1996) documented interannual shifts in pollinator dominance, with bats prevailing in one year and hummingbirds in other, a pattern linked to El Niño events that alter rainfall, vegetation, and bat abundance. We propose that *H. acranthus* may follow a similar strategy, with mixed floral traits enabling dual pollination. Such flexibility is likely advantageous in the seasonal lomas ecosystem, where extreme climatic events like El Niño strongly influence resource availability (Ferreyra 1993; Cano *et al*. 1999). The hypothesized migratory behavior of *P. genovensium* (Sahley 1996; Sahley & Baraybar 1996; Ossa *et al*. 2020) further underscores the value of maintaining alternative pollinators: hummingbirds may provide reliable service during periods of reduced bat activity, thereby supporting reproductive success under fluctuating conditions (Muchhala & Thomson 2010). However, as this study was conducted over a single year, our ability to characterize temporal fluctuations in pollinator presence and confirm seasonal patterns remains limited. Finally, the disjunct distribution of *H. acranthus* along the Peruvian coast, with populations separated by large distances, aligns with bat-mediated pollination, as bats are highly effective long-distance pollen vectors, promoting genetic connectivity despite habitat fragmentation (Fleming *et al*. 2009).

### Mixed-pollination strategies in a broader context

The most effective pollinator principle states that floral traits should evolve toward the optimum imposed by the pollinator contributing the most to reproductive success (Stebbins 1970), but this is not always observed. When no single guild imposes consistent selection, phenotypes compatible with multiple guilds can be favored because their combined contributions exceed those provided by specialization: a mixed strategy is more convenient when pollinator abundances shift across space or time (Kay & Anderson, 2025).

Mixed pollination can occur even in species with clear floral syndromes. In *Marginatocereus marginatus*, competition with co-flowering columnar cacti reduced the reliability of specific pollinator guilds, and dual nocturnal-diurnal anthesis allowed reliance on both bats and hummingbirds (Dar et al. 2006). We consider it unlikely that *H. pseudomelanostele* or *L. acanthurus* generate meaningful pollination interference for *H. acranthus* at our site, as both species occur at much lower densities, and *L. acanthurus* shows little to no flowering overlap with *H. acranthus*. However, this is a possibility in other lomas populations of coastal Peru, where this cactus coexists with multiple hummingbird- and bat-pollinated species, often showing reproductive compatibility even across genera (Arakaki et al. 2021).

Geographic variation in pollination systems has been documented in cacti. Schlumpberger et al. (2009) found shifts from bee pollination (short, morning-opening, low-nectar flowers) sphingid pollination (long-tubed, dusk-opening, nectar-rich flowers) across populations of *Echinopsis ancistrophora* along an altitudinal gradient in Argentina. In Mexico, *Stenocereus thurberi* also showed spatial differences: northern and central populations relied mainly on bats with no pollen limitation, whereas southern populations on mixed pollinators and showed pollen limitation, likely due to temporal variation in pollinator availability (Bustamante et al. 2010). *Pachycereus pecten-aboriginum* also displayed geographic differentiation in anthesis and nectar secretion that aligns with nocturnal versus mixed nocturnal-diurnal pollination in Valiente-Banuet et al. (2004). This tendency to more generalized pollination systems has also been documented in other Pachycereeae such as *Carnegia gigantea* (Fleming et al., 1996); supporting that generalized strategies appear to be more reliable in zones where bats predictability is lower (e.g., northern Mexico versus central and Southern regions, Rojas-Martínez et al. 1999). These dynamics remain unstudied in Peruvian cacti. The mixed pollination strategy we propose for *H. acranthus* may be shaped by local factors such as habitat condition, pollinator abundance, and the presence or absence of co-flowering cacti. However, since our data comes from a single site and one sampling year, it cannot reveal if pollination mechanisms vary across space or time, or whether such variation explains possible differences among populations. Wider geographic coverage and long-term studies will be essential to address these questions.

## Conclusions

Our study shows that *H. acranthus* exhibits floral traits primarily aligned with bat pollination while remaining compatible with diurnal hummingbird pollination, with both groups showing behavior consistent with effective pollen transfer. Bees and sphingids mismatched key floral traits and likely acted primarily as nectar or pollen thieves. We found no differences in floral traits or visitation frequencies between color morphs, providing no support for pollinator-driven divergence between morphotypes. Overall, our findings indicate a mixed vertebrate pollination system in which reliance on multiple guilds may enhance reproductive stability in the strongly seasonal lomas ecosystem, where pollinator availability can shift across space and time, maintaining interactions with multiple pollinator guilds may enhance reproductive stability. This study represents one of the few contributions to understanding cactus pollination in this biogeographically singular environment, and highlights the need for broader spatial and temporal studies across Peruvian cacti communities.

## Acknowledgements

We thank A. Ceroni and M. Flores (Weberbauer Herbarium-MOL) for their support in obtaining funding and contributions during the design of the study; C. Reynel and S. Terreros (Forestry Herbarium-MOLF) for botanical samples reception; M. Alvarado and E. Medina (Department of Entomology-UNMSM) for arthropod samples reception and assitance on taxonomic identification. This research was funded by the ‘XI Concurso de fondos de investigación para círculos de investigación 2021’ from UNALM (Lima, Peru). This study complies with legal requirements set by MINAGRI and SERFOR for specimen collection (RD N° D0000046-2024-MIDAGRI-SERFOR-DGGSPFFS-DGSPF).

