## Supplementary Figures for "Mixed-vertebrate pollination traits and pollinators of *Haageocereus acranthus* (Cactaceae) in a lomas desert ecosystem of coastal Peru"

**Figure S1.** Principal Component Analysis (PCA) of floral morphology for white and pink-red morphotypes of *Haageocereus acranthus*. Panels show the distribution of individuals along (A) PC1 (39.64%) and PC2 (24.67%), and (B) PC1 (39.64%) and PC3 (11.57%). Arrows indicate the direction and relative contribution of each floral trait to the principal components. Points represent individual flowers, colored by morphotype (green = white, red = pink-red).

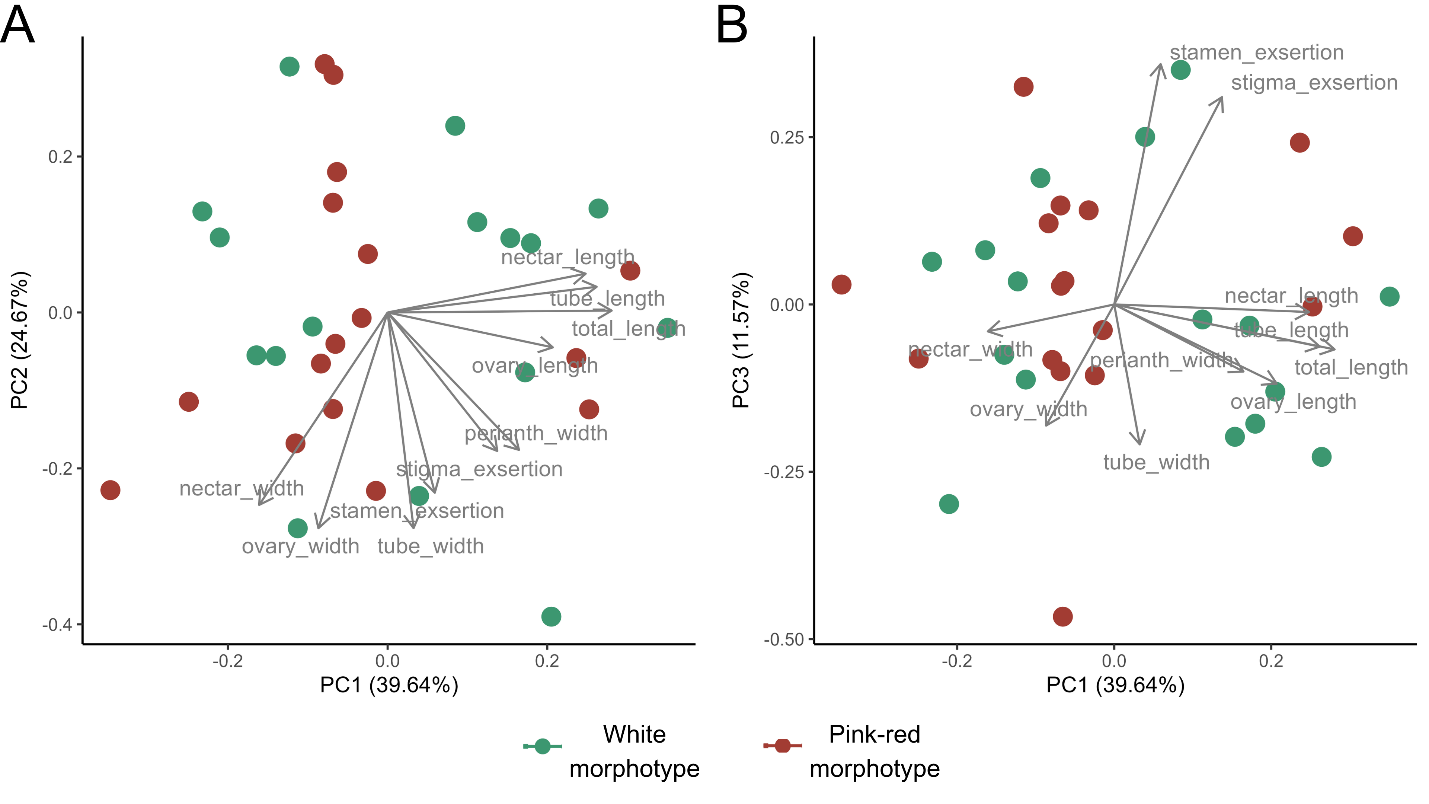

**Figure S2.** Model diagnostic plots generated using the DHARMa package for the selected flower aperture model. (A) QQ plot of scaled residuals. No significant deviations detected in the Kolmogorov-Smirnov test (p = 0.930), dispersion test (p = 0.888), or outlier test (p = 1). (B) Residuals vs. predicted plot indicates significant quantile deviations, however, this pattern was observed across all candidate models, suggesting it reflects an inherent characteristic of the data, more than model misspecification. Zero-inflation test resulted significant (p < 0.001), however, all zero values present in the dataset correspond to biologically expected observations.

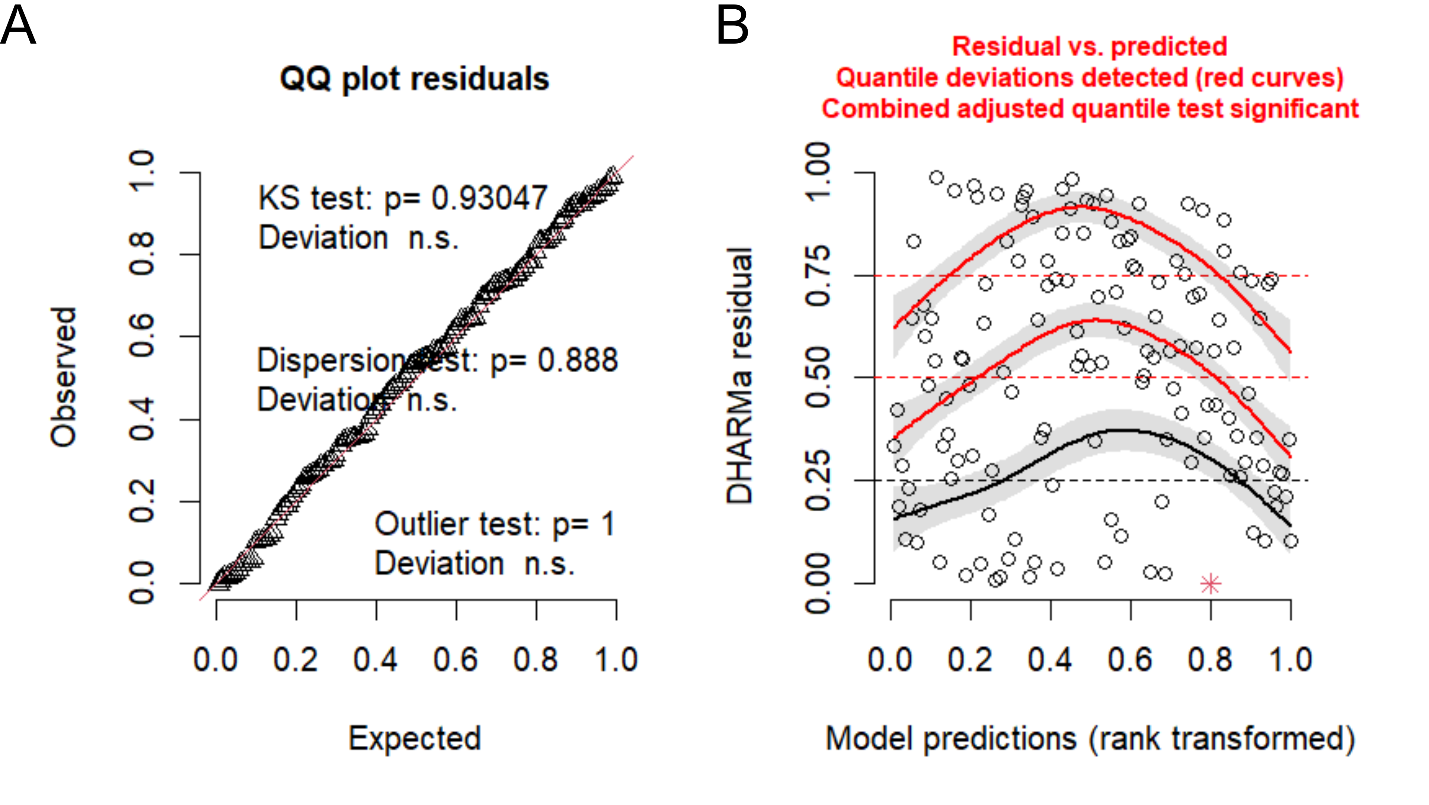

**Figure S3.** Bootstrap distributions of fixed effect estimates for the selected flower aperture model, based on 1000 cluster bootstrap iterations. Each histogram shows the distribution of a given coefficient across bootstrap resamples, with the original model estimate indicated by the red vertical line and the 95% bootstrap confidence interval by the blue dashed vertical lines. Bootstrapped estimates were stable and well-centered within confidence intervals across all model terms.

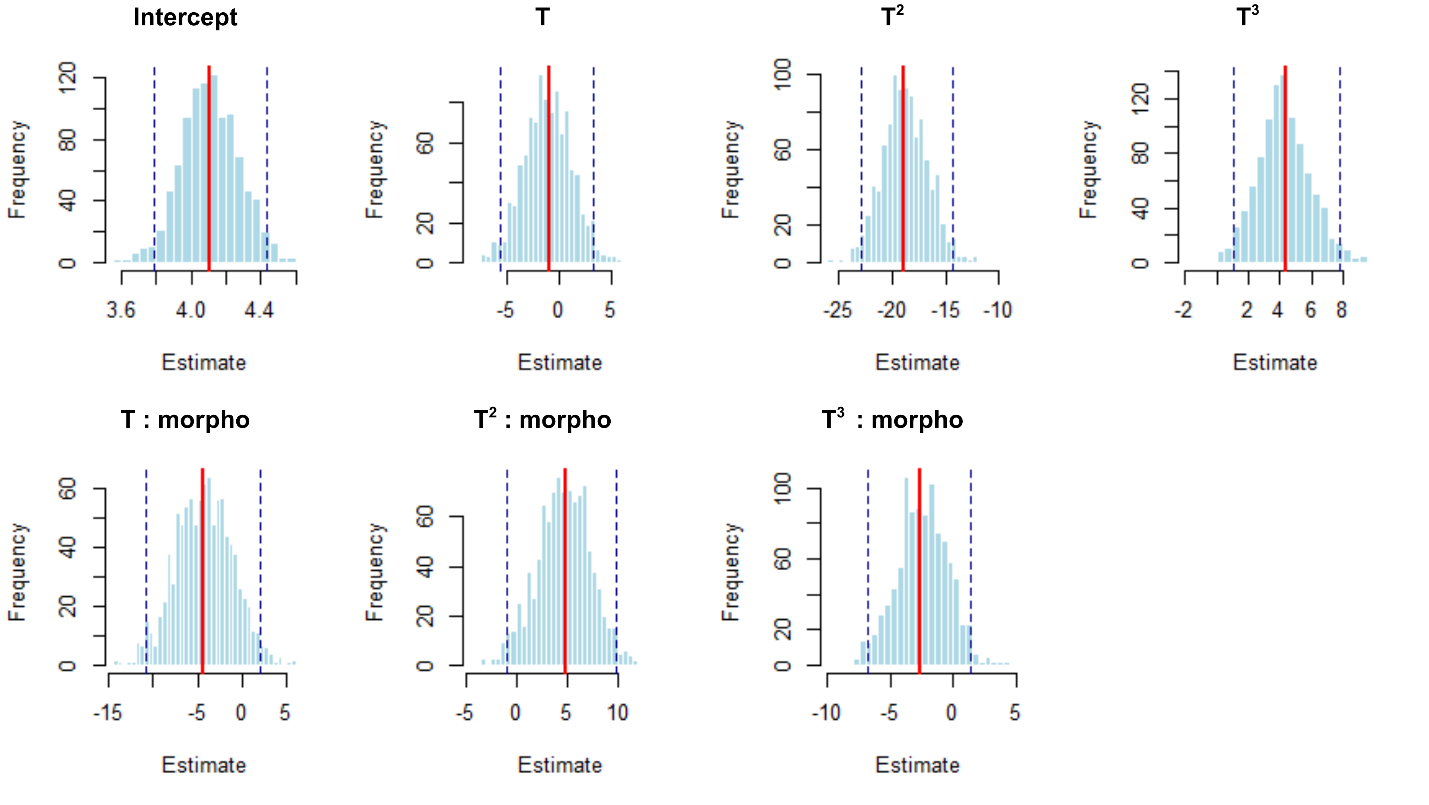

**Figure S4.** Model diagnostic plots generated using the DHARMa package for the selected nectar volume model. (A) QQ plot of scaled residuals. No significant deviations detected in the Kolmogorov-Smirnov test (p = 0.093) nor dispersion test (p = 0.696); although the outlier test was significant (p = 0.019), the flagged values were biologically reasonable and therefore retained. (B) Residuals vs. predicted plot indicated non significant quantile deviations. The zero-inflation test resulted non significant (p = 0.92).

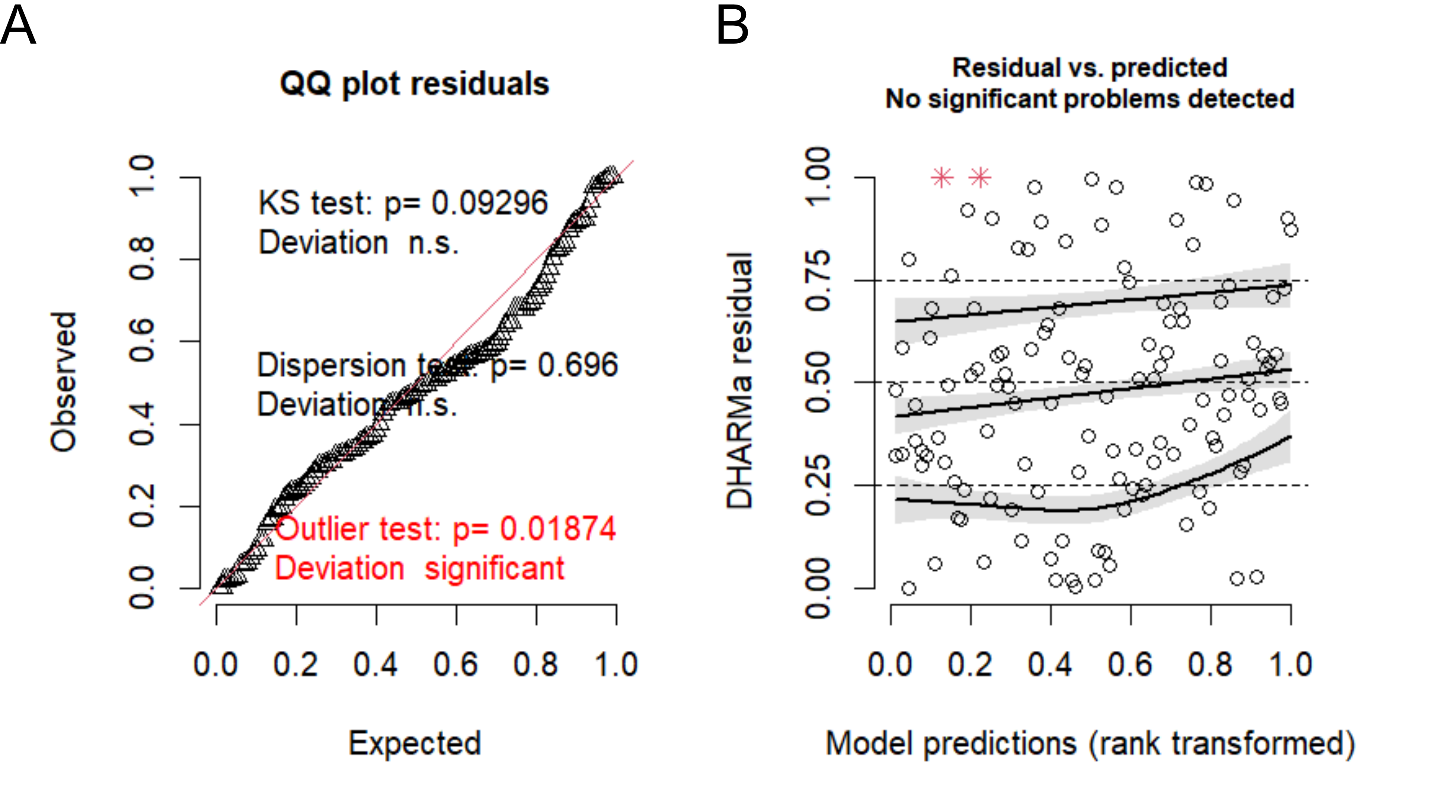

**Figure S5.** Bootstrap distributions of fixed effect estimates for the selected nectar volume model, based on 1000 cluster bootstrap iterations. Each histogram shows the distribution of a given coefficient across bootstrap resamples, with the original model estimate indicated by the red vertical line and the 95% bootstrap confidence interval by the blue dashed vertical lines. Bootstrapped estimates were stable and well-centered within confidence intervals across all model terms.

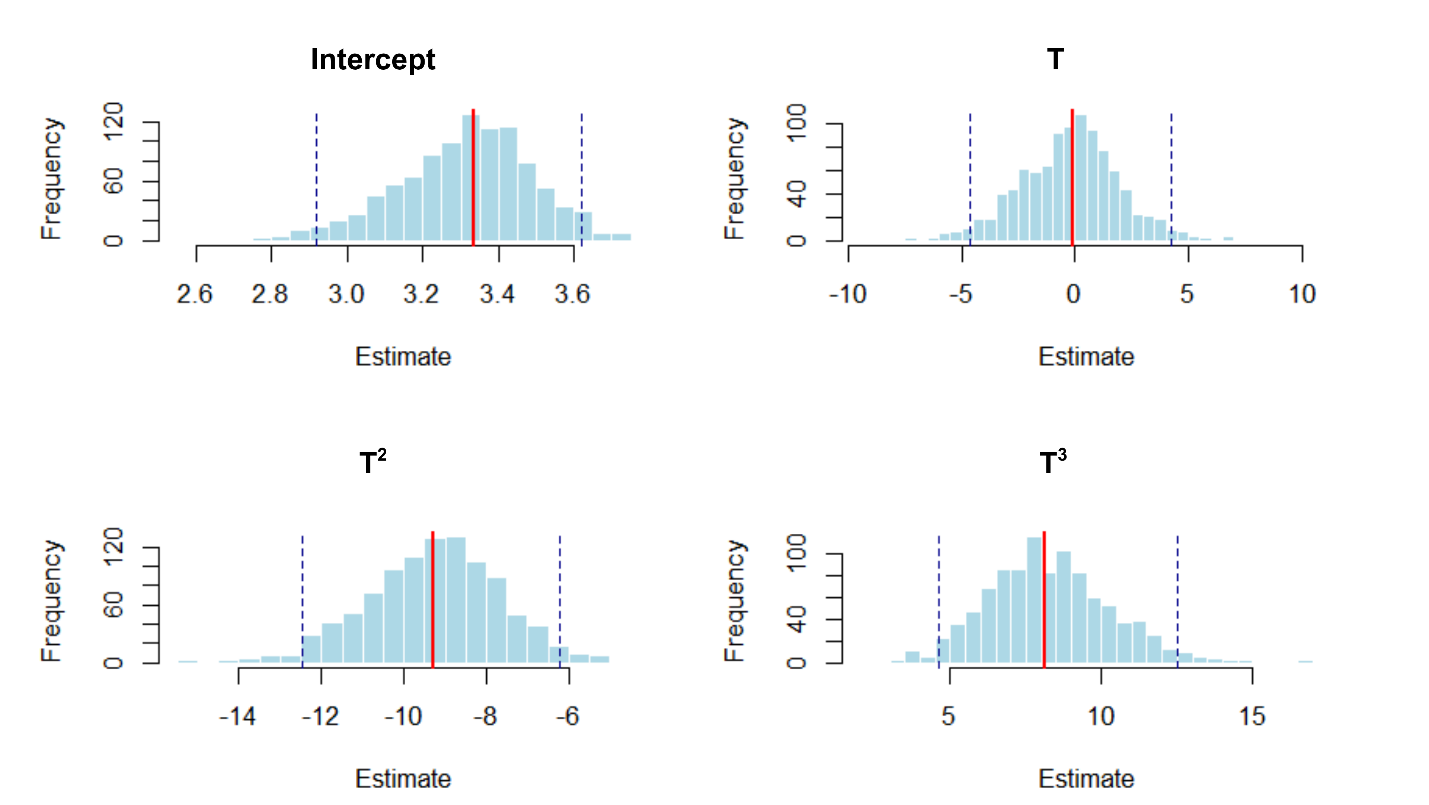

**Figure S6.** Model diagnostic plots generated using the DHARMa package for the selected nectar sugar concentration model. (A) QQ plot of scaled residuals. No significant deviations detected in the Kolmogorov-Smirnov test (p = 0.905), dispersion test (p = 0.709), or outlier test (p = 1). (B) Residuals vs. predicted plot indicated non significant quantile deviations. The zero-inflation test resulted non significant (p = 1).

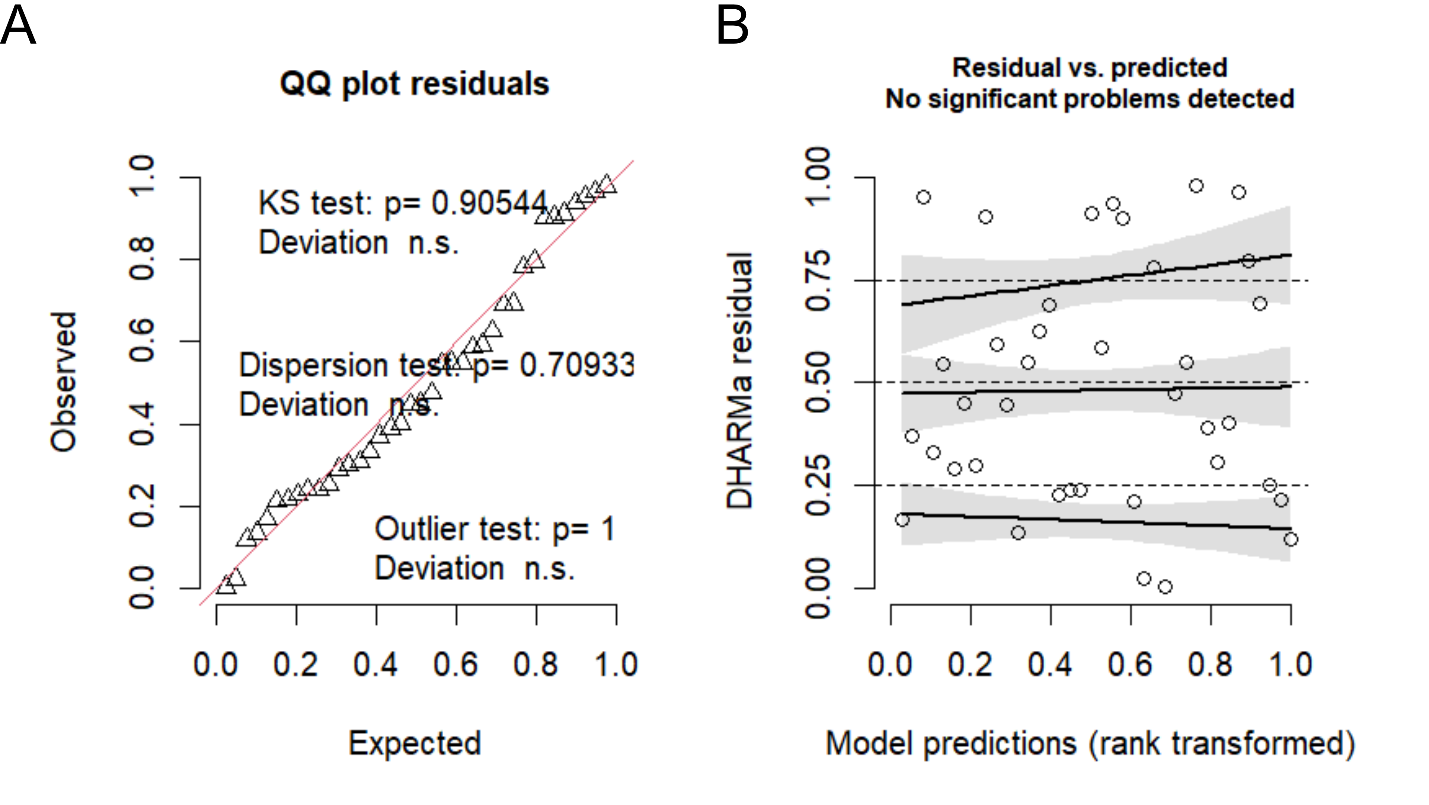

**Figure S7.** Bootstrap distributions of fixed effect estimates for the selected nectar sugar concentration model, based on 1000 cluster bootstrap iterations. Each histogram shows the distribution of a given coefficient across bootstrap resamples, with the original model estimate indicated by the red vertical line and the 95% bootstrap confidence interval by the blue dashed vertical lines. Bootstrapped estimates were stable and centered within confidence intervals across all model terms.

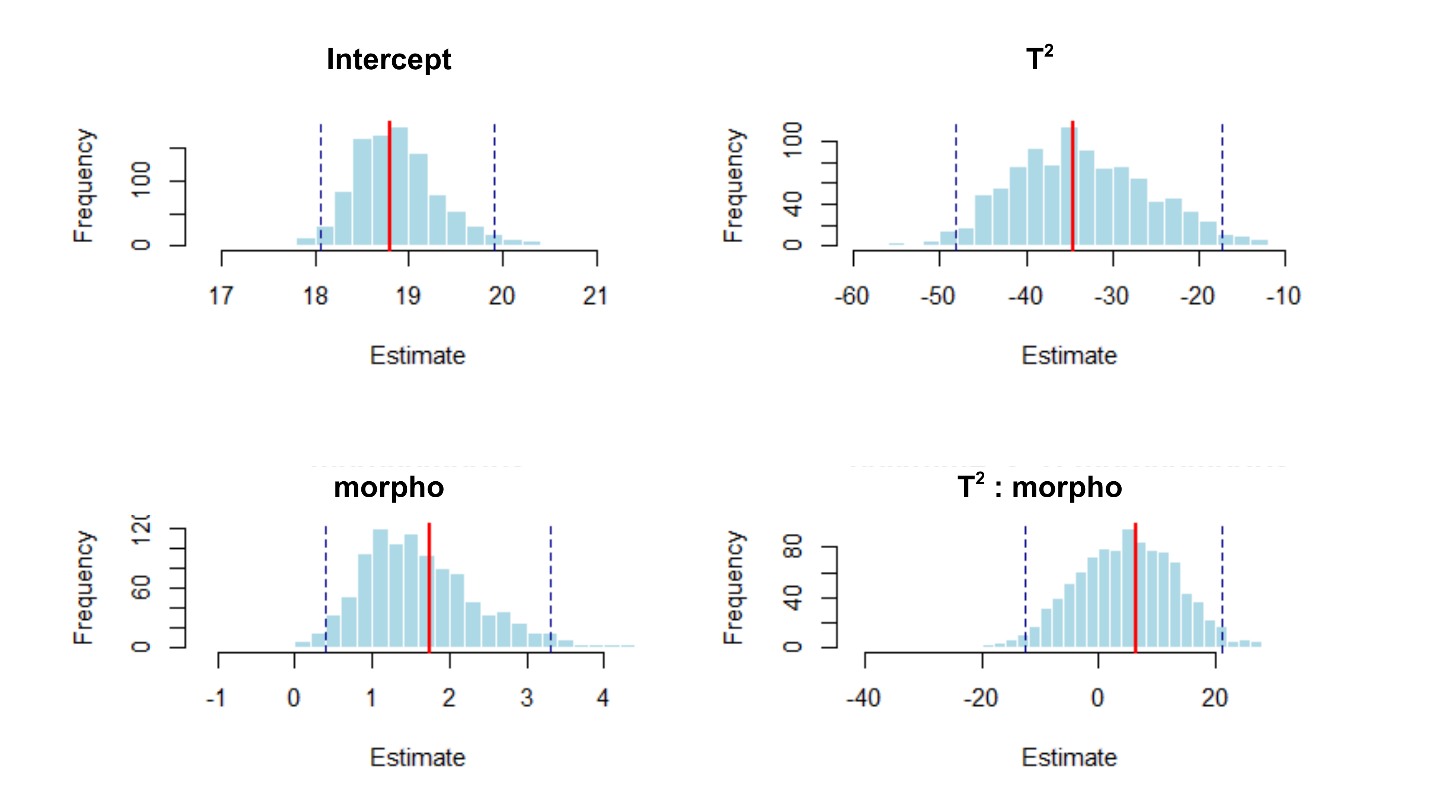

**Figure S8.** Model diagnostic plots generated using the DHARMa package for the selected nectar energy content model. (A) QQ plot of scaled residuals. No significant deviations detected in the Kolmogorov-Smirnov test (p = 0.792), dispersion test (p = 0.992), or outlier test (p = 0.391). (B) Residuals vs. predicted plot indicated significant quantile deviations, however, this pattern was observed across all candidate models. The zero-inflation test resulted non significant (p = 1).

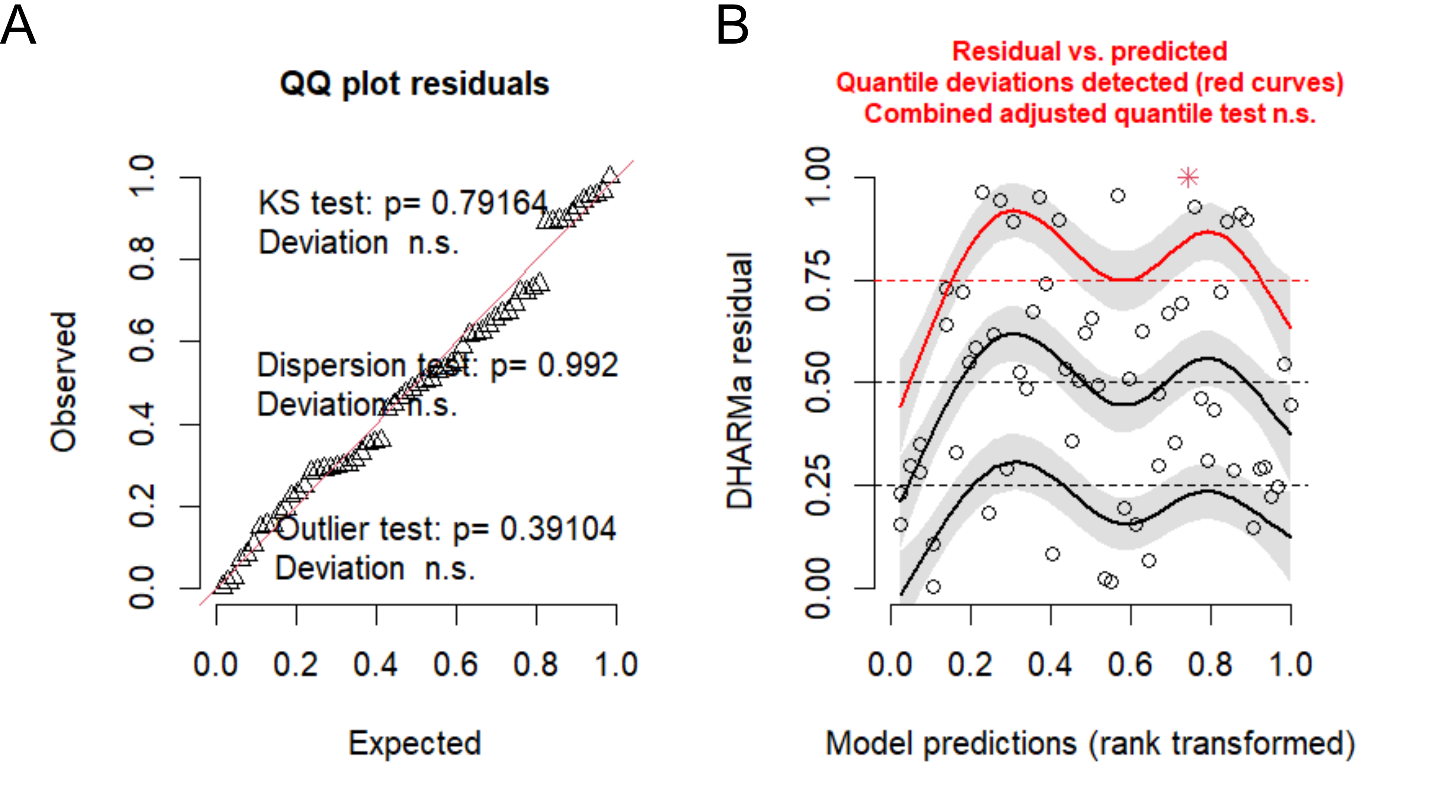

**Figure S9.** Bootstrap distributions of fixed effect estimates for the selected nectar energy content model, based on 1000 cluster bootstrap iterations. Each histogram shows the distribution of a given coefficient across bootstrap resamples, with the original model estimate indicated by the red vertical line and the 95% bootstrap confidence interval by the blue dashed vertical lines. Bootstrapped estimates were stable and well-centered within confidence intervals across all model terms.

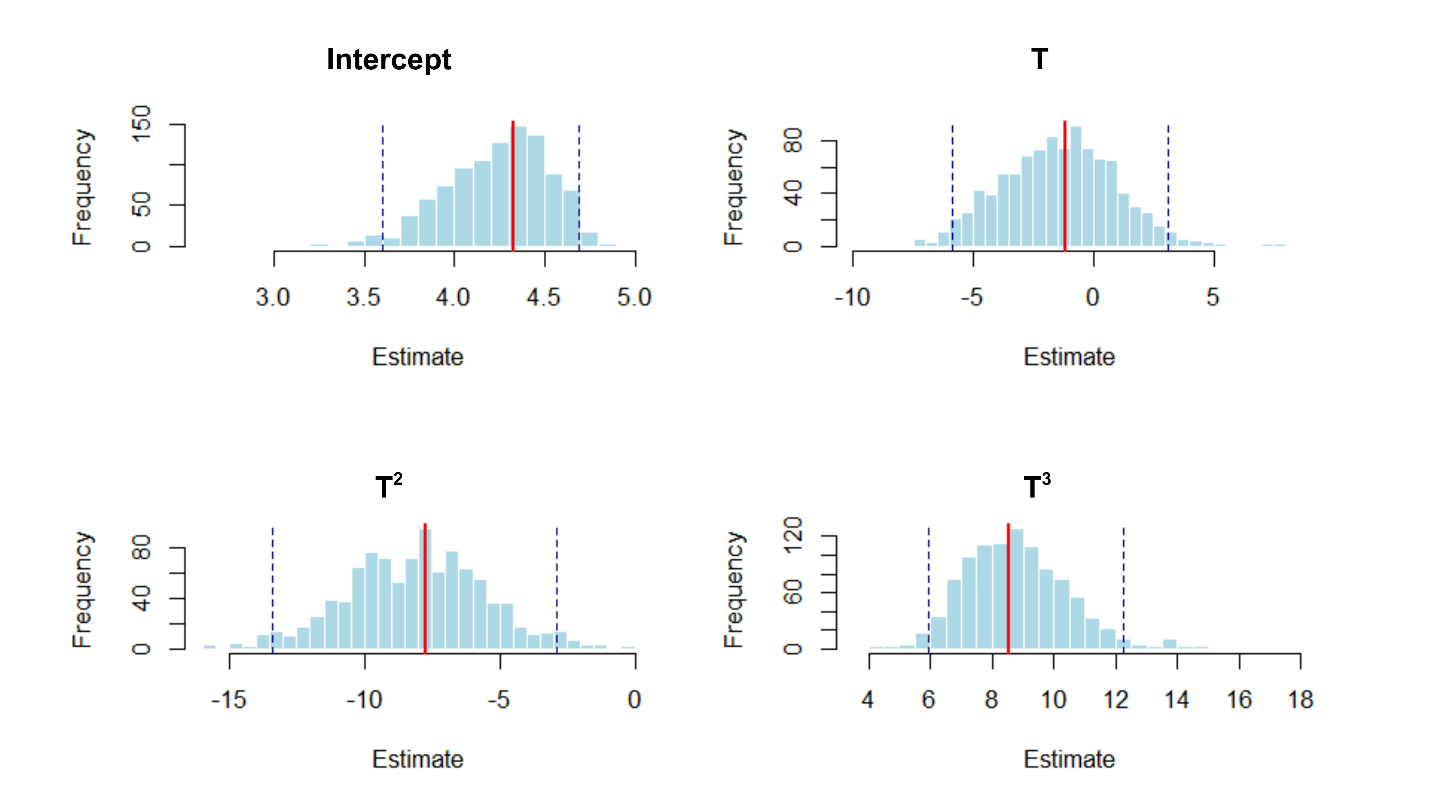

**Figure S10.** Floral visitors of Haageocereus acranthus: (A) Linepithema sp. (Formicidae); (B–C) Solenopsis sp. (Formicidae); (D) Carpophilus sp. (Nitidulidae); (E–F) Anyphaenidae.

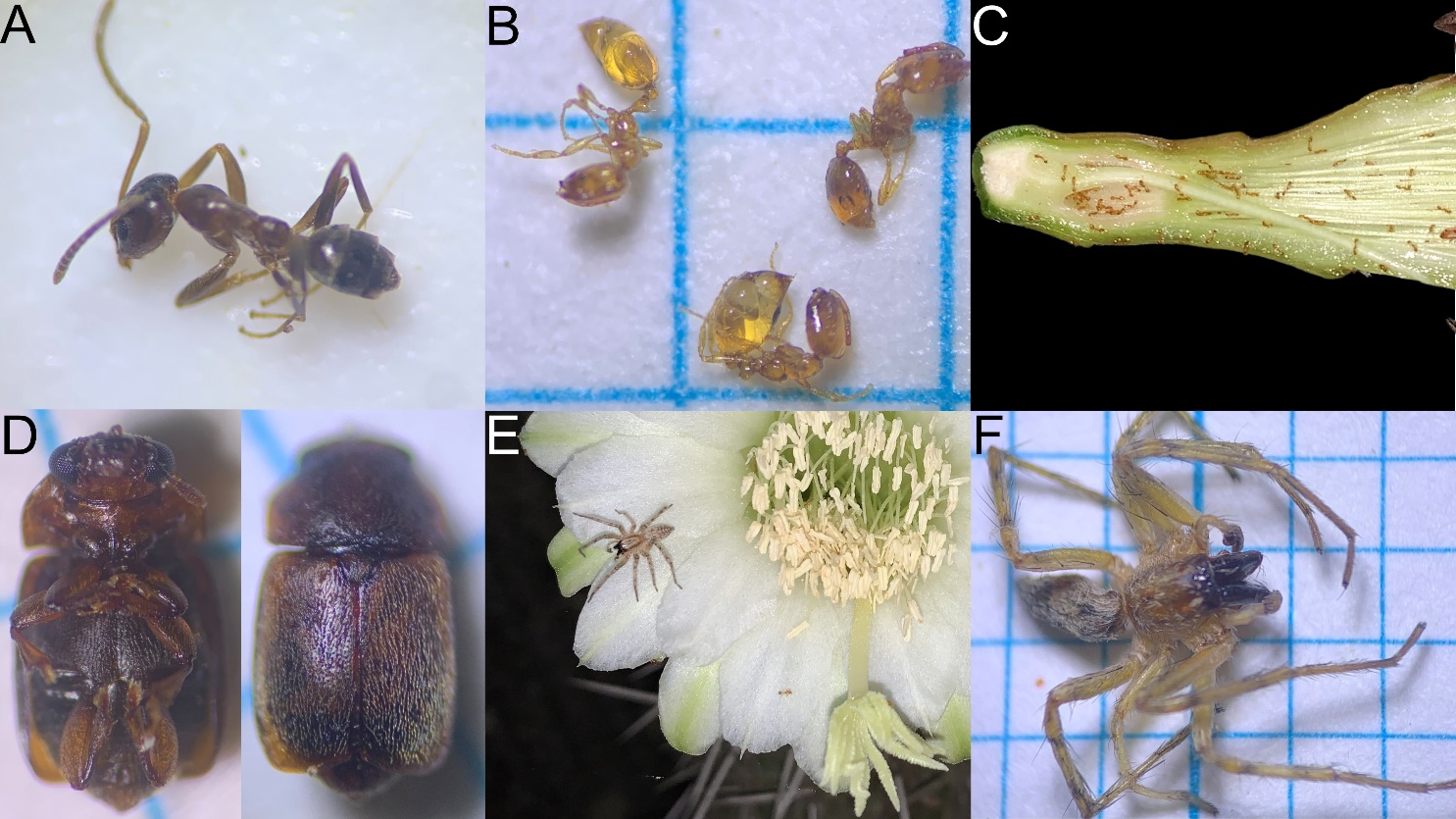

**Figure S11.** Floral visitors of *H. acranthus*: (A) Thomisidae*;* (B) Salticidae; (C) Phoridae; (E) *Apis mellifera* (Apidae); (F) Halictidae.

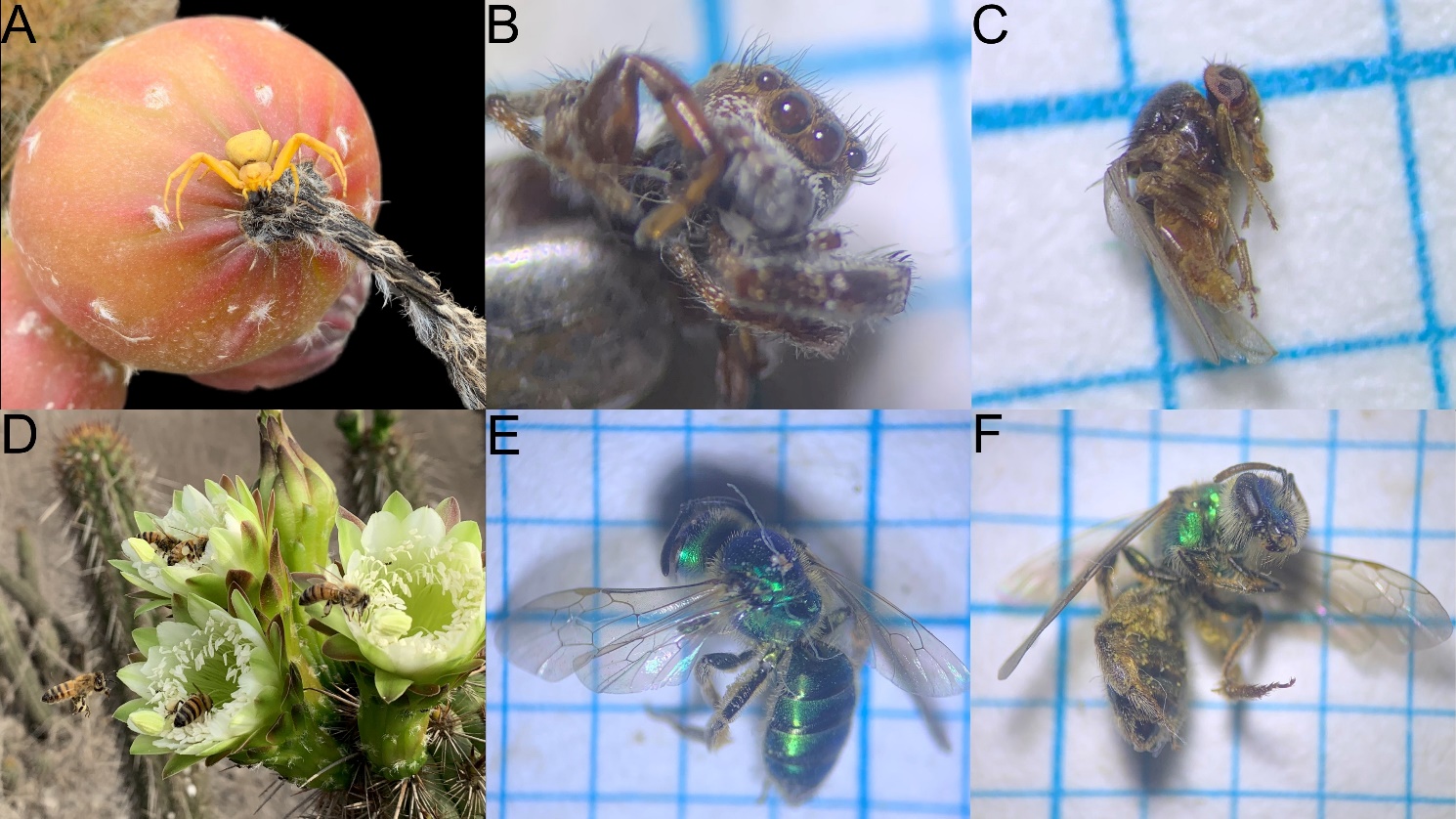

**Figure S12.** Model diagnostic plots generated using the DHARMa package for the selected flower visit model. (A) QQ plot of scaled residuals. No significant deviations detected in the Kolmogorov-Smirnov test (p = 0.311), dispersion test (p = 0.688), or outlier test (p = 1). (B) Residuals vs. predicted plot indicated non significant quantile deviations. The zero-inflation test resulted non significant (p = 0.424).

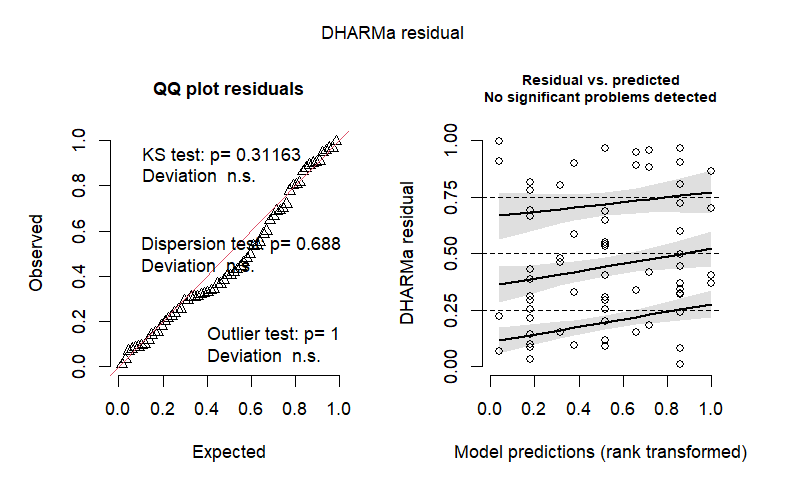

**Figure S13.** Bootstrap distributions of fixed effect estimates for the selected flower visitation model, based on 1000 cluster bootstrap iterations. Each histogram shows the distribution of a given coefficient across bootstrap resamples, with the original model estimate indicated by the red vertical line and the 95% bootstrap confidence interval by the blue dashed vertical lines. Bootstrap distributions were highly concentrated around the original model estimates, with the vast majority of iterations producing stable estimates consistent with the reported coefficients. However, 95% bootstrap confidence intervals were wide due to a small proportion of resamples producing extreme estimates, likely reflecting occasional resamples where visitor representation was severely imbalanced across flowers.

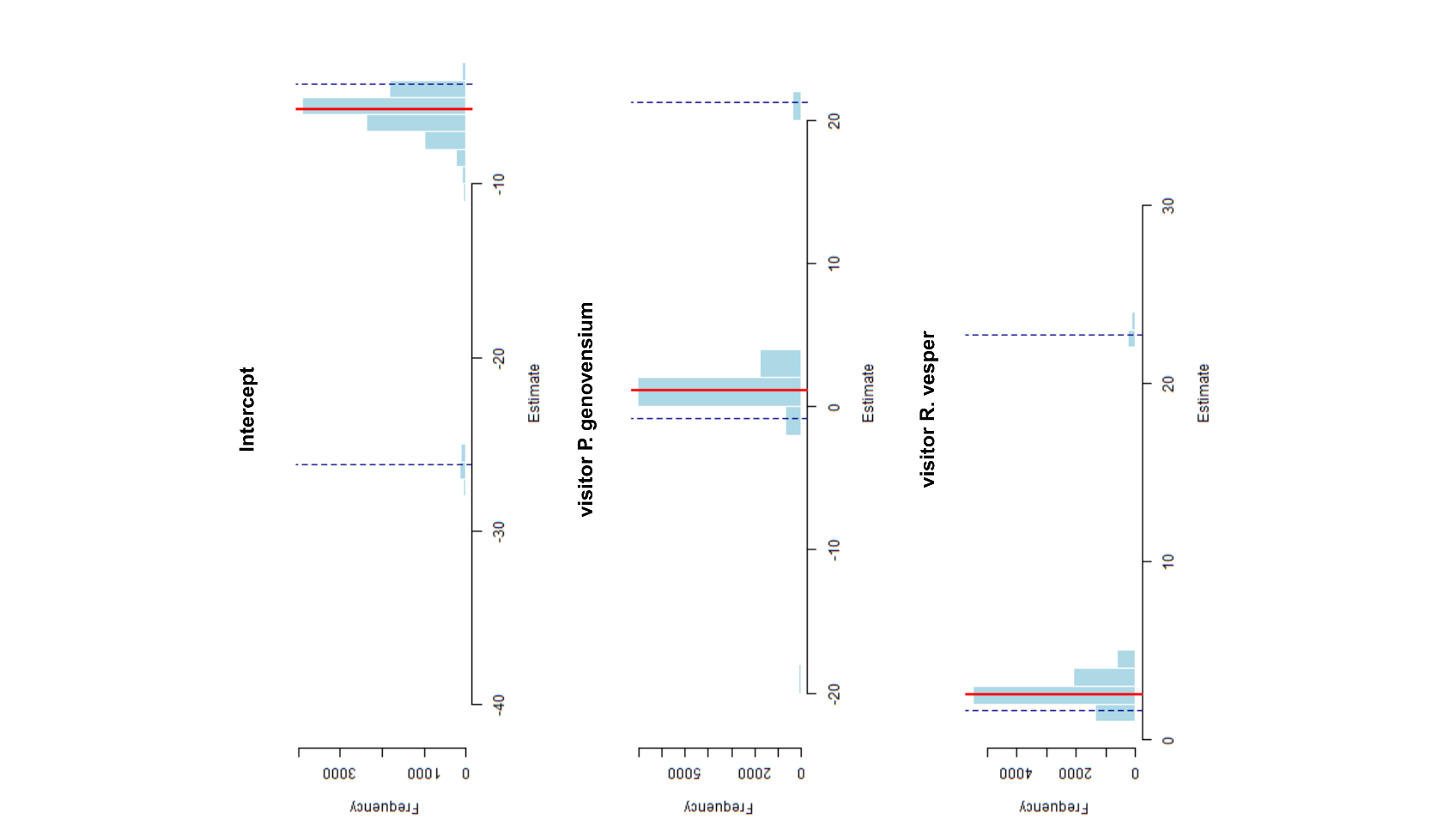

**Figure S14.** Air temperature through time recorded through 2022 by the VILLA MARIA DEL TRIUNFO meteorological station (code 112233; 12°9'59'' S, 76°55'12'' W; 292 m.a.s.l.), Lima, Peru. This station is located 8.5 km from the study area and is the nearest meteorological station available. Blue dots indicate hourly air temperature measurements (°C), while the red line represents a LOESS-smoothed trend. Data were provided by the Peruvian National Meteorology and Hydrology Service (SENAMHI). Relative humidity data were not available for this station during the study period.

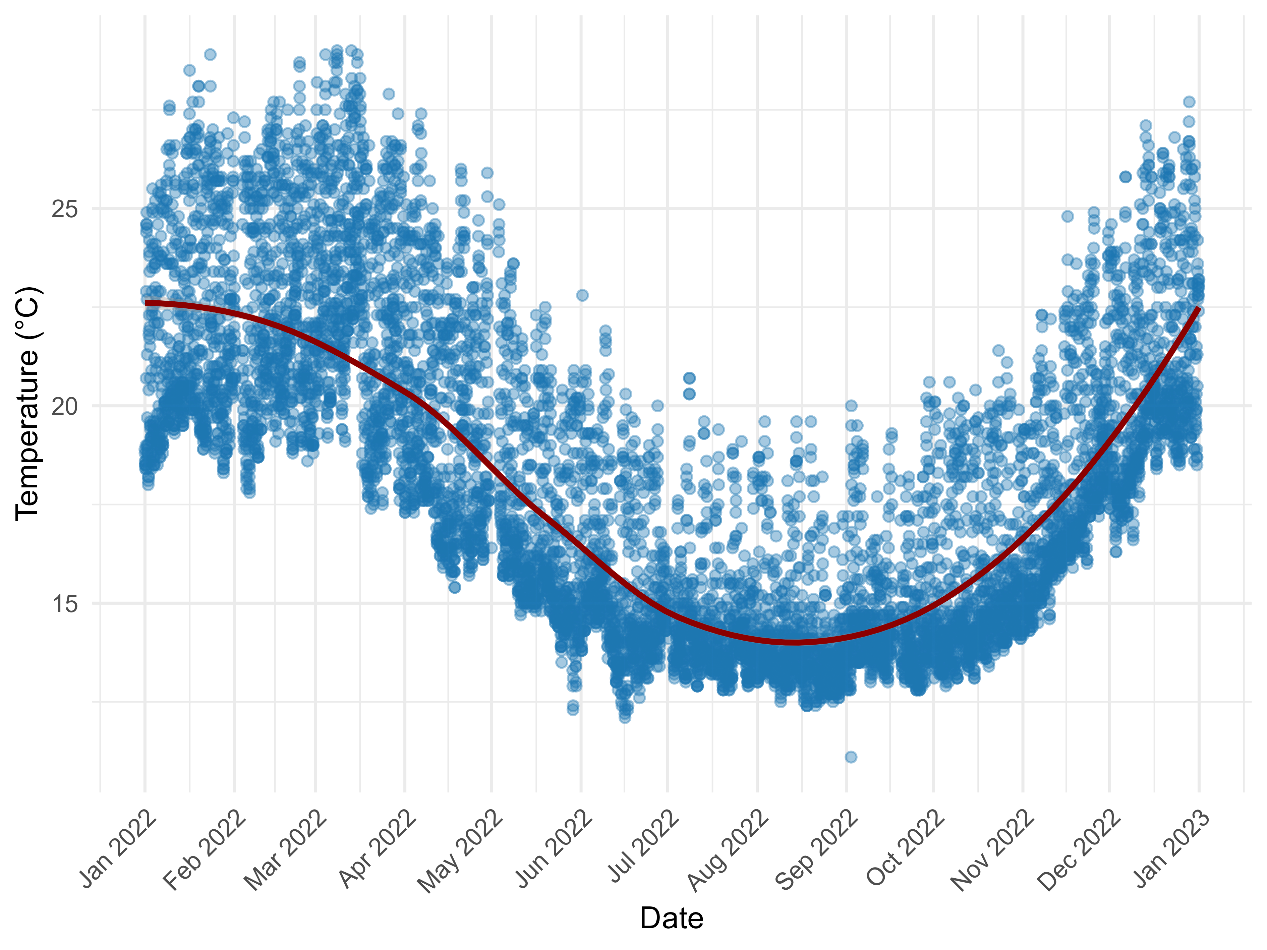
