## Supplementary Tables for "Mixed-vertebrate pollination traits and pollinators of *Haageocereus acranthus* (Cactaceae) in a lomas desert ecosystem of coastal Peru"

**Table S1.** Annual phenology of *H. acranthus* in 2022. Values represent the mean ± standard error (SE) of the number of buds, flowers, unripe fruits, and ripe fruits recorded per month.

| **Month** | **Mean ± SE** | | | | | | | | | | | |
| --- | --- | --- | --- | --- | --- | --- | --- | --- | --- | --- | --- | --- |
|  | **Buds** | | | **Flowers** | | | **Unripe fruits** | | | **Ripe fruits** | | |
| Jan | 4.50 | ± | 1.33 | 2.07 | ± | 0.40 | 2.77 | ± | 0.68 | 1.00 | ± | 0.46 |
| Feb | 5.60 | ± | 1.17 | 3.53 | ± | 0.82 | 3.23 | ± | 0.93 | 0.60 | ± | 0.27 |
| Mar | 0.73 | ± | 0.23 | 1.03 | ± | 0.39 | 1.57 | ± | 0.43 | 1.90 | ± | 0.70 |
| Apr | 0.93 | ± | 0.39 | 1.20 | ± | 0.53 | 0.73 | ± | 0.45 | 0.70 | ± | 0.22 |
| May | 0.87 | ± | 0.39 | 0.10 | ± | 0.07 | 0.57 | ± | 0.30 | 0.10 | ± | 0.07 |
| Jul | 0.72 | ± | 0.29 | 0.07 | ± | 0.07 | 0.20 | ± | 0.20 | 0.13 | ± | 0.10 |
| Sep | 0.87 | ± | 0.39 | 0.10 | ± | 0.06 | 0.07 | ± | 0.07 | 0.03 | ± | 0.03 |
| Oct | 1.47 | ± | 0.34 | 0.27 | ± | 0.11 | 0.17 | ± | 0.08 | 0.00 | ± | 0.00 |
| Nov | 7.97 | ± | 1.92 | 1.83 | ± | 0.71 | 3.23 | ± | 0.90 | 0.37 | ± | 0.20 |
| Dec | 7.30 | ± | 1.85 | 4.10 | ± | 0.87 | 5.93 | ± | 1.24 | 0.63 | ± | 0.28 |

**Table S2.** Morphometric measurements of floral traits for white and pink-red floral morphs of ***H. acranthus****.* Values represent mean ± standard error (SE) of length in cm.

| **Trait** | **Mean ± SE** | | | | | |
| --- | --- | --- | --- | --- | --- | --- |
|  | **White morph** | | | **Pink-red morph** | | |
| Total length | 6.47 | ± | 0.28 | 6.18 | ± | 0.27 |
| Tube length | 4.88 | ± | 0.24 | 4.73 | ± | 0.21 |
| Perianth width | 6.39 | ± | 0.20 | 6.20 | ± | 0.18 |
| Tube width | 1.99 | ± | 0.07 | 1.96 | ± | 0.06 |
| Stigma exsertion | 3.50 | ± | 0.16 | 3.36 | ± | 0.12 |
| Stamen exsertion | 2.01 | ± | 0.09 | 2.11 | ± | 0.11 |
| Ovary length | 0.87 | ± | 0.06 | 0.76 | ± | 0.04 |
| Ovary width | 0.74 | ± | 0.04 | 0.75 | ± | 0.03 |
| Nectar chamber length | 1.56 | ± | 0.10 | 1.53 | ± | 0.09 |
| Nectar chamber width | 0.72 | ± | 0.04 | 0.80 | ± | 0.05 |

**Table S3.** Model selection results for flower aperture. The table presents the fixed effects included in each model and their respective model performance metrics: degrees of freedom (df), log-likelihood (logLik), corrected Akaike Information Criterion (AICc), delta AICc (ΔAICc), and model weight. M represents the flower morphotype, and T is the time after midday in its linear (T), quadratic (T²), and cubic (T³) forms. Interaction terms (M : T, M : T², M : T³) are also included. Models are ranked by AICc, with lower values indicating better fit.

| **Fixed effects** | | | | | | | **df** | **logLik** | **AICc** | **delta** | **weight** |
| --- | --- | --- | --- | --- | --- | --- | --- | --- | --- | --- | --- |
| **M** | **T** | **M : T** | **T^2^** | **M : T^2^** | **T^3^** | **M : T^3^** |  |  |  |  |  |
|  | + | + | + | + | + | + | 9 | -223.95 | 467.29 | 0.00 | 0.560 |
| + | + | + | + | + | + | + | 10 | -223.94 | 469.60 | 2.31 | 0.176 |
|  | + |  | + |  | + |  | 6 | -228.55 | 469.74 | 2.45 | 0.165 |
|  | + | + | + | + |  |  | 7 | -228.35 | 471.56 | 4.27 | 0.066 |
| + | + | + | + | + |  |  | 8 | -228.35 | 473.81 | 6.52 | 0.022 |
|  | + |  | + |  |  |  | 5 | -232.38 | 475.21 | 7.92 | 0.011 |
|  | + |  |  |  |  |  | 4 | -289.03 | 586.36 | 119.07 | 0.000 |
|  | + | + |  |  |  |  | 5 | -288.03 | 586.51 | 119.22 | 0.000 |
| + | + | + |  |  |  |  | 6 | -287.99 | 588.61 | 121.32 | 0.000 |

**Table S4.** Model selection results for nectar volume. The table presents the fixed effects included in each model and their respective model performance metrics: degrees of freedom (df), log-likelihood (logLik), corrected Akaike Information Criterion (AICc), delta AICc (ΔAICc), and model weight. M represents the flower morphotype, and T is the time after midday in its linear (T), quadratic (T²), and cubic (T³) forms. Interaction terms (M : T, M : T², M : T³) are also included. Models are ranked by AICc, with lower values indicating better fit.

| **Fixed effects** | | | | | | | **df** | **logLik** | **AICc** | **delta** | **weight** |
| --- | --- | --- | --- | --- | --- | --- | --- | --- | --- | --- | --- |
| **M** | **T** | **M : T** | **T^2^** | **M : T^2^** | **T3** | **M : T^3^** |  |  |  |  |  |
|  | + |  | + |  | + |  | 7 | -505.35 | 1025.65 | 0.00 | 0.920 |
|  | + | + | + | + | + | + | 10 | -504.60 | 1031.11 | 5.46 | 0.060 |
| + | + | + | + | + | + | + | 11 | -504.52 | 1033.35 | 7.71 | 0.020 |
|  | + |  | + |  |  |  | 6 | -516.14 | 1044.99 | 19.34 | 0.000 |
|  | + | + | + | + |  |  | 8 | -515.61 | 1048.46 | 22.81 | 0.000 |
| + | + | + | + | + |  |  | 9 | -515.54 | 1050.63 | 24.98 | 0.000 |
|  | + |  |  |  |  |  | 5 | -532.72 | 1075.94 | 50.29 | 0.000 |
|  | + | + |  |  |  |  | 6 | -532.37 | 1077.44 | 51.80 | 0.000 |
| + | + | + |  |  |  |  | 7 | -532.35 | 1079.66 | 54.01 | 0.000 |

**Table S5.** Model selection results for nectar sugar concentration. The table presents the fixed effects included in each model and their respective model performance metrics: degrees of freedom (df), log-likelihood (logLik), corrected Akaike Information Criterion (AICc), delta AICc (ΔAICc), and model weight. M represents the flower morphotype, and T is the time after midday in its linear (T), quadratic (T²), and cubic (T³) forms. Interaction terms (M : T, M : T², M : T³) are also included. Models are ranked by AICc, with lower values indicating better fit.

| **Fixed effects** | | | | | | | **df** | **logLik** | **AICc** | **delta** | **weight** |
| --- | --- | --- | --- | --- | --- | --- | --- | --- | --- | --- | --- |
| **M** | **T** | **M : T** | **T^2^** | **M : T^2^** | **T^3^** | **M : T^3^** |  |  |  |  |  |
|  | + |  | + |  | + |  | 6 | -74.29 | 163.30 | 0.00 | 0.374 |
|  | + |  |  |  |  |  | 4 | -77.39 | 164.00 | 0.72 | 0.261 |
| + | + | + |  |  |  |  | 6 | -75.18 | 165.10 | 1.80 | 0.152 |
|  | + |  | + |  |  |  | 5 | -77.25 | 166.40 | 3.09 | 0.080 |
|  | + | + |  |  |  |  | 5 | -77.38 | 166.60 | 3.36 | 0.070 |
| + | + | + | + | + |  |  | 8 | -73.77 | 168.51 | 5.23 | 0.027 |
|  | + | + | + | + |  |  | 7 | -75.81 | 169.40 | 6.08 | 0.018 |
| + | + | + | + | + | + | + | 10 | -71.11 | 170.40 | 7.08 | 0.011 |
|  | + | + | + | + | + | + | 9 | -73.54 | 171.50 | 8.22 | 0.006 |

**Table S6.** Model selection results for nectar energy content. The table presents the fixed effects included in each model and their respective model performance metrics: degrees of freedom (df), log-likelihood (logLik), corrected Akaike Information Criterion (AICc), delta AICc (ΔAICc), and model weight. M represents the flower morphotype, and T is the time after midday in its linear (T), quadratic (T²), and cubic (T³) forms. Interaction terms (M : T, M : T², M : T³) are also included. Models are ranked by AICc, with lower values indicating better fit.

| **Fixed effects** | | | | | | | **df** | **logLik** | **AICc** | **delta** | **weight** |
| --- | --- | --- | --- | --- | --- | --- | --- | --- | --- | --- | --- |
| **M** | **T** | **M : T** | **T^2^** | **M : T^2^** | **T^3^** | **M : T^3^** |  |  |  |  |  |
|  | + |  | + |  | + |  | 7 | -261.99 | 540.06 | 0 | 0.738 |
|  | + | + | + | + | + | + | 10 | -259.4 | 543.11 | 3.0518 | 0.160 |
| + | + | + | + | + | + | + | 11 | -258.37 | 544.01 | 3.9571 | 0.102 |
| + | + | + | + | + |  |  | 9 | -269.59 | 560.64 | 20.579 | 0.000 |
|  | + |  | + |  |  |  | 6 | -273.95 | 561.43 | 21.37 | 0.000 |
|  | + | + | + | + |  |  | 8 | -272.97 | 564.66 | 24.604 | 0.000 |
|  | + |  |  |  |  |  | 5 | -287.24 | 585.54 | 45.486 | 0.000 |
|  | + | + |  |  |  |  | 6 | -287.16 | 587.85 | 47.797 | 0.000 |
| + | + | + |  |  |  |  | 7 | -286.22 | 588.52 | 48.461 | 0.000 |

**Table S7.** Total number of visits and mean number of visits per hour recorded for each floral visitor for each *Haageocereus* morphotype (White and Red-pink) individually and pooled across both morphotypes. Visitors include sphingids, *Rhodopis vesper*, and *Platalina genovensium*.

| **Visitor** | **Morphotype** | **Number of visits** | **Mean number of visits per hour** |
| --- | --- | --- | --- |
| Sphingids | White | 5 | 0.0303 |
|  | Red-pink | 1 | 0.0065 |
|  | Pooled | 6 | 0.184 |
| R. vesper | White | 13 | 0.0677 |
|  | Red-pink | 7 | 0.0303 |
|  | Pooled | 20 | 0.049 |
| P. genovensium | White | 45 | 0.254 |
|  | Red-pink | 35 | 0.187 |
|  | Pooled | 80 | 0.22 |

**Table S8.** Model selection results for visitation frequency. The table presents the fixed effects included in each model and their respective model performance metrics: degrees of freedom (df), log-likelihood (logLik), corrected Akaike Information Criterion (AICc), delta AICc (ΔAICc), and model weight. M represents the flower morphotype, V the flower visitor, and M : V the interaction between both fixed effects. Models are ranked by AICc, with lower values indicating better fit.

| **Fixed effects** | | | **df** | **logLik** | **AICc** | **delta** | **weight** |
| --- | --- | --- | --- | --- | --- | --- | --- |
| **V** | **M** | **M : V** |  |  |  |  |  |
| + |  |  | 4 | -100.46 | 209.58 | 0.00 | 0.717 |
| + | + |  | 5 | -100.45 | 211.89 | 2.32 | 0.225 |
| + | + | + | 7 | -99.35 | 214.63 | 5.05 | 0.057 |
